# Rosetta FunFolDes – a general framework for the computational design of functional proteins

**DOI:** 10.1101/378976

**Authors:** Jaume Bonet, Sarah Wehrle, Karen Schriever, Che Yang, Anne Billet, Fabian Sesterhenn, Andreas Scheck, Freyr Sverrisson, Sabrina Vollers, Roxanne Lourman, Melanie Villard, Stéphane Rosset, Bruno E. Correia

## Abstract

The robust computational design of functional proteins has the potential to deeply impact translational research and broaden our understanding of the determinants of protein function, nevertheless, it remains a challenge for state-of-the-art methodologies. Here, we present a computational design approach that couples conformational folding with sequence design to embed functional motifs into heterologous proteins. We performed extensive benchmarks, where the most unexpected finding was that the design of function into proteins may not necessarily reside in the global minimum of the energetic landscape, which could have important implications in the field. We have computationally designed and experimentally characterized a distant structural template and a *de novo* “functionless” fold, two prototypical design challenges, to present important viral epitopes. Overall, we present an accessible strategy to repurpose old protein folds for new functions, which may lead to important improvements on the computational design of functional proteins.

## Introduction

Proteins are one of the main functional building blocks of the cell. The ability to create novel proteins outside of the natural realm has opened the path towards innovative achievements, such as new protein pathways (Cross et al., 2017), cellular functions (Joh et al., 2014), and therapeutic leads (Correia et al., 2010; Correia et al., 2014; Kulkarni et al., 2015). Computational protein design is the rational and structure-based approach to solve the inverse folding problem, i.e. the search for the best putative sequence capable of fitting and stabilizing a given protein’s three-dimensional conformation (Coluzza, 2017). As such, a great deal of effort has been placed into the understanding of the rules of protein folding and stability (Koga et al., 2012; Marcos et al., 2017) and its relation to the appropriate sequence space (Kuhlman & Baker, 2000).

Computational protein design focus on two main axes of search related to the structural and sequence spaces that are explored. Fixed backbone approaches work with a static protein backbone conformation, which greatly constrains the sequence space that is explored by the computational algorithm (Kuhlman & Baker, 2000). Following the same principles as naturally occurring homologs, which often exhibit a certain degree of structural diversity, flexible backbone approaches enhance the sequence diversity, adding the challenge of identifying energetically favorable sequence variants that are correctly coupled to the structural perturbations (Murphy et al., 2012).

Another variation for computational design approaches is *de novo* design, in which protein backbones are assembled *in silico*, followed by sequence optimization to fold into a pre-defined three-dimensional conformation without being constrained by previous sequence information (Hill, Raleigh, Lombardi, & DeGrado, 2000). This approach tests our understanding of the rules governing the structure of different protein folds. The failures and successes of this approach confirm and correct the principles used for the protein design process (Koga et al., 2012; Marcos et al., 2017).

One of the main aims of computational protein design is the rational design of functional proteins capable of carrying existing or novel functions into new structural contexts (Street & Mayo, 1999). One can broadly classify three main approaches for the design of functional proteins: redesigning of pre-existing functions, grafting of functional sites onto heterologous proteins, and designing of novel functions not found in the protein repertoire. The redesign of a preexisting function to alter its catalytic activity (Yu et al., 2014) or improve its binding target recognition (Guntas, Purbeck, & Kuhlman, 2010) can be considered the most conservative approach; as it is typically accomplished by point mutations around the functional area of interest, it tends to have little impact on global structure and stability of the designed protein. On the other hand, the design of fully novel functions has most noticeably been achieved by applying chemical principles that tested our fundamental knowledge of enzyme catalysis (Jiang et al., 2008; Kries, Blomberg, & Hilvert, 2013).

Between these two approaches resides protein grafting. This method aims to repurpose natural folds as carriers for exogenous known functions. It relies on the strong relationship between protein structure and activity, to allocate a given functionality from one protein to another by means of transferring the structural motif responsible for the function (Azoitei et al., 2011; Correia et al., 2011; Correia et al., 2010; Correia et al., 2014; Kulkarni et al., 2015; Procko et al., 2014; Viana et al., 2013). The most successful grafting approaches are highly dependent on structural similarity between the functional motif and the insertion region in the protein scaffold. When the functional motif and the insertion region are almost identical in backbone conformation, functional transfer can be performed by side-chain grafting, i.e. mutating the target residues into those of the functional motif (Correia et al., 2010; Kulkarni et al., 2015). In much more challenging scenarios, full backbone grafting may be used in conjunction with directed evolution to make the structure fully compatible with the new function (Azoitei et al., 2011). Nevertheless, motif transfer is limited between very similar structural regions, which greatly constrains the subset of putative scaffolds that can be used for this purpose, especially as the structural complexity of the functional motif grows.

Previously, we have demonstrated the possibility of expanding protein grafting to scaffolds with segments that have low structural similarity. To accomplish that task, we developed a prototype protocol named Rosetta Fold From Loops (FFL) (Correia et al., 2014; Procko et al., 2014).

The distinctive feature of our protocol is the coupling of the folding and design stages to bias the sampling towards structural conformations and sequences that stabilize the grafted functional motif. In the past, FFL was used to obtain designs that were functional (synthetic immunogens (Correia et al., 2014) and protein-based inhibitors (Procko et al., 2014)) and where the experimentally determined crystal structures closely resembled the computational models; however, the structures of the functional sites were still very close to the insertion segments of the hosting scaffolds.

Here, we present a complete re-implementation of the FFL protocol with enhanced functionalities, simplified user interface and complete integration with any other available Rosetta protocols. We have called this new, more generalist protocol Rosetta Functional Folding and Design (FunFolDes), we have benchmarked it in a number of scenarios providing important technical details to better exploit and expand the capabilities of the original protocol. Furthermore, we challenged FunFolDes with two design tasks to probe the boundaries of applicability of the protocol. The design tasks were centered on using distant structural template as hosting scaffold and functionalizing a *de novo* designed protein - in both challenges, FunFolDes succeeded in functionalizing the designed proteins. These results are encouraging and provide a solid basis for the broad applicability of FunFolDes as a strategy for the robust computational design of functionalized proteins.

## Results

### Rosetta FunFolDes - a computational framework for design of functional proteins

The original prototype of the Rosetta Fold From Loops (FFL) protocol was successfully used to transplant the structural motif of the Respiratory Syncytial Virus (RSV) protein F site II neutralizing epitope into a protein scaffold in the context of a vaccine design application (Correia et al., 2014).

FFL enabled the insertion and conformational stabilization of the structural motif into a defined protein topology by using Rosetta’s fragment insertion machinery to fold the polypeptide chain to adopt the desired topology (Rohl, Strauss, Misura, & Baker, 2004) which was then sequence designed. Information content from the scaffold structure was used to guide the folding, ensuring an overall similar topology while allowing for the conformational changes needed to stabilize the inserted structural motif.

The final implementation of the protocol, referred to as FunFolDes, is schematically represented in **Figure 1**, and fully described in Materials and Methods. Our upgrades to FFL focused on three main aims: I) improve the applicability of the system to allow handling of more complex structural motifs; II) enhance the design of functional proteins by including binding partners in the simulations; III) offer a higher degree of control over each stage of the simulation while improving the usability for non-experts. These three aims were achieved through the implementation of five core technical improvements described below.

**Figure 1.**
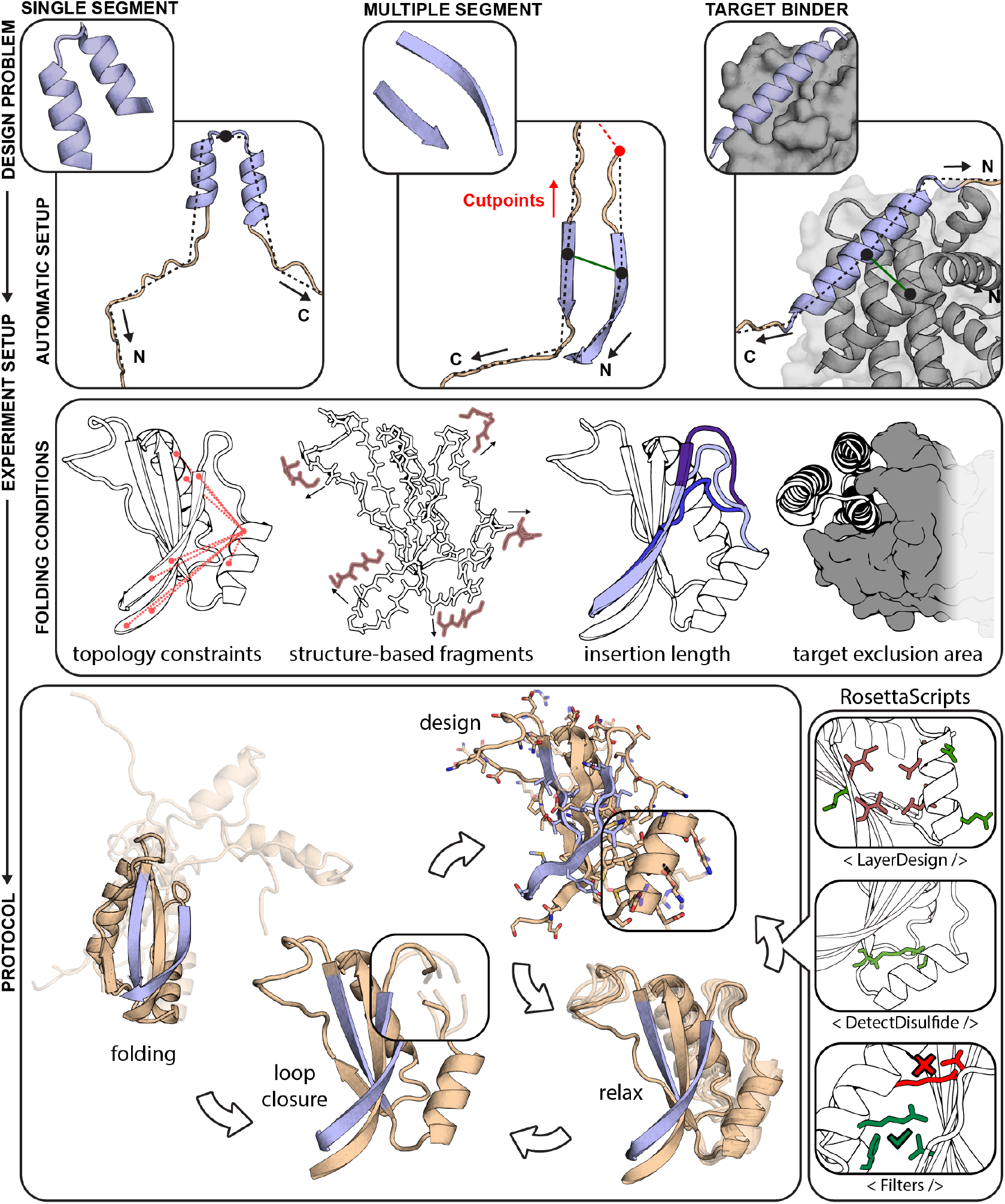
Rosetta FunFolDes - method overview. FunFolDes was devised to tackle a wide range of functional protein design problems, combining a higher user control of the simulation parameters whilst simultaneously lowering the level of expertise required. FunFolDes is able to transfer single- and multi-segment motifs together with the target partner by exploiting Rosetta’s FoldTree framework (top row). A wider range of information can be extracted from the template to shift the final conformation towards a more productive design space (middle row), including targeted distance constraints, generation of structure-based fragments, motif insertion in sites with different residue length and presence of the binding target to bias the folding stage. The bottom row showcases the most typical application of the FunFolDes protocol. Integration in RosettaScripts allows to tailor FunFolDes behavior and for a seamless integration with other protocols, and complex selection logics can be added to address the different complexities in each design task.

#### Insertion of multi-segment functional sites

The initial implementation of FFL was limited to the insertion of a single-segment structural motif, which was sufficient to demonstrate its potential at the time (Correia et al., 2014; Procko et al., 2014). However, most functional sites in proteins typically entail, at the structural level, multiple discontinuous segments; which is the case for protein-protein interfaces or enzyme active-sites, among others (Aragues, Sali, Bonet, Marti-Renom, & Oliva, 2007; Richter, Leaver-Fay, Khare, Bjelic, & Baker, 2011). FunFolDes can now handle functional sites with any number of discontinuous segments, ensuring the native orientations of each of the segments. Furthermore, it allows control of the backbone flexibility of each of the insertion points and the order in the protein scaffold sequence in which each segment is inserted. Finally, the sequence length between the motif and the insertion region is not required to be the same, allowing the user to search for protein scaffolds using alternative metrics to the full backbone RMSDs between the motif and the protein scaffold (Azoitei et al., 2011; Correia et al., 2010). These new features essentially allow the replacement between completely different structural segments. Thus, they greatly enhance the types of structural motifs that can be targeted with FunFolDes, widening the applicability of the computational protocol.

#### Structural folding and sequence design in the presence of a binding partner

Many of the functional roles of proteins in cells require physical interaction with other proteins, nucleic acids, or metabolites (Garcia-Garcia et al., 2012). Several proposed mechanisms to regulate binding affinities and specificities in protein interactions involve protein flexibility, such as induced fit and conformational selection (Chakrabarti et al., 2016; Lange et al., 2008). Inspired by these naturally occurring mechanisms, we devised a strategy to fold and design in the presence of the desired binding partner. Including the binder in simulations has a twofold benefit. On the one hand, is a way of explicitly representing functional constraints to bias the designed protein towards a functional sequence space, resolving putative clashes derived from the template scaffold and, thus, significantly enlarging the number of usable templates. On the other hand, this approach facilitates the design of new additional contact residues (outside of the motif) that may afford enhanced affinity and/or specificity. Here, we tested FunFolDes in a model system for which extensive experimental data has been collected, and we show how this approach improves the sampling of productive conformational and sequence space.

#### Region-specific structural constraints

FFL could exploit distance constraints from the target scaffold to guide the folding stage. A simplified solution was implemented in FFL with two possible simulation modes, where either constraints are collected throughout the protein scaffold, or folding is unconstrained. Currently, FunFolDes can collect from full-template to region-specific constraints, allowing greater levels of flexibility in areas of the scaffold that can be critical for function (e.g. segments close to the interface of a target protein) and improving the sampling of conformations which otherwise could be missed or highly underrepresented. Furthermore, FunFolDes is no longer limited to atom-pair distance constraints (Rohl & Baker, 2002) and can incorporate other types of kinematic constraints, such as angle and dihedral constraints (Bowers, Strauss, & Baker, 2000), which have been used to improve success rates when folding scaffolds rich in beta-strands (Marcos et al., 2017).

#### On-the-fly fragment picking

Fragment insertion is a core algorithm in Rosetta protocols exploring high degrees of freedom of the polypeptide chain, such as *ab initio* protein prediction (Simons, Ruczinski, et al., 1999), loop modeling (Stein & Kortemme, 2013), or more recently, FFL (Correia et al., 2014). Classically, fragment libraries are generated through sequence-based predictions of secondary structure and dihedral angles (Bowers et al., 2000). This information is used in a Rosetta application to obtain three- and nine residue-long fragment libraries from naturally occurring proteins, which are then provided to the downstream protocols. Leveraging internal functionalities in Rosetta, FunFolDes can assemble fragment sets automatically. Due to its particularities, secondary structure, dihedral angles, and accessible solvent area can be automatically computed from the protein scaffold’s structure. Although sequence-based fragments can still be provided, this removes the need for secondary applications in the protocol pipeline, boosting the usability of FunFolDes by lowering the barrier for non-experts. It also enables the assembly of protocols in which the fragment sets are mutable along the procedure. The benchmark presented in this paper evaluates the performance of such functionality.

#### Compatibility with other Rosetta modules

Finally, FunFolDes is compatible with Rosetta’s modular xml-interface: Rosetta Scripts (RS) (Fleishman et al., 2011). This enables customization of the FunFolDes protocol and, more importantly, connection with other protocols and filters available through the RS interface. In order to obtain a full integration with this interface, the FunFolDes protocol is divided in multiple *Movers* (i.e. modules capable of altering the information content of a structure).

We devised two benchmark scenarios to test the performance of FunFolDes. One of these aimed to capture conformational changes in small protein domains caused by sequence insertions or deletions, and the second scenario assessed protocol performance to fold and design a binder in the presence of the target binding partner.

### Capturing conformational and sequence changes in small protein domains

Typical protein design benchmarks are assembled by stripping native side chains from known protein structures and evaluating the sequence recovery of the design algorithm (Kuhlman & Baker, 2000). The main design aim of FunFolDes is to insert structural motifs into protein folds while allowing flexibility across the overall structure. This conformational freedom allows the full protein scaffold to adapt and stabilize the functional motif’s conformation. This is a main distinctive point from other approaches to design functional proteins that rely on a mostly rigid scaffold (Azoitei et al., 2011; Correia et al., 2010; Fallas et al., 2017; Hill et al., 2000; Joh et al., 2014; Richter et al., 2011). For many modeling problems, such as protein structure prediction, protein-protein and protein-ligand docking, or protein design, standardized benchmark datasets are available (Vreven et al., 2015) or easily accessible. Devising a benchmark for designed proteins with propagating conformational changes across the structure is challenging, as we are assessing both structural accuracy as well as sequence recovery of the protocol.

To address this problem, we analyzed structural domains found repeatedly in natural proteins and clustered them according to their definition in the CATH database (Dawson et al., 2017). As a result, we were able to select a set of 14 benchmark targets labeled T01 through T14 (**Figure 2A**). A detailed description on the construction of the benchmark can be found in the Materials and Methods section.

**Figure 2.**
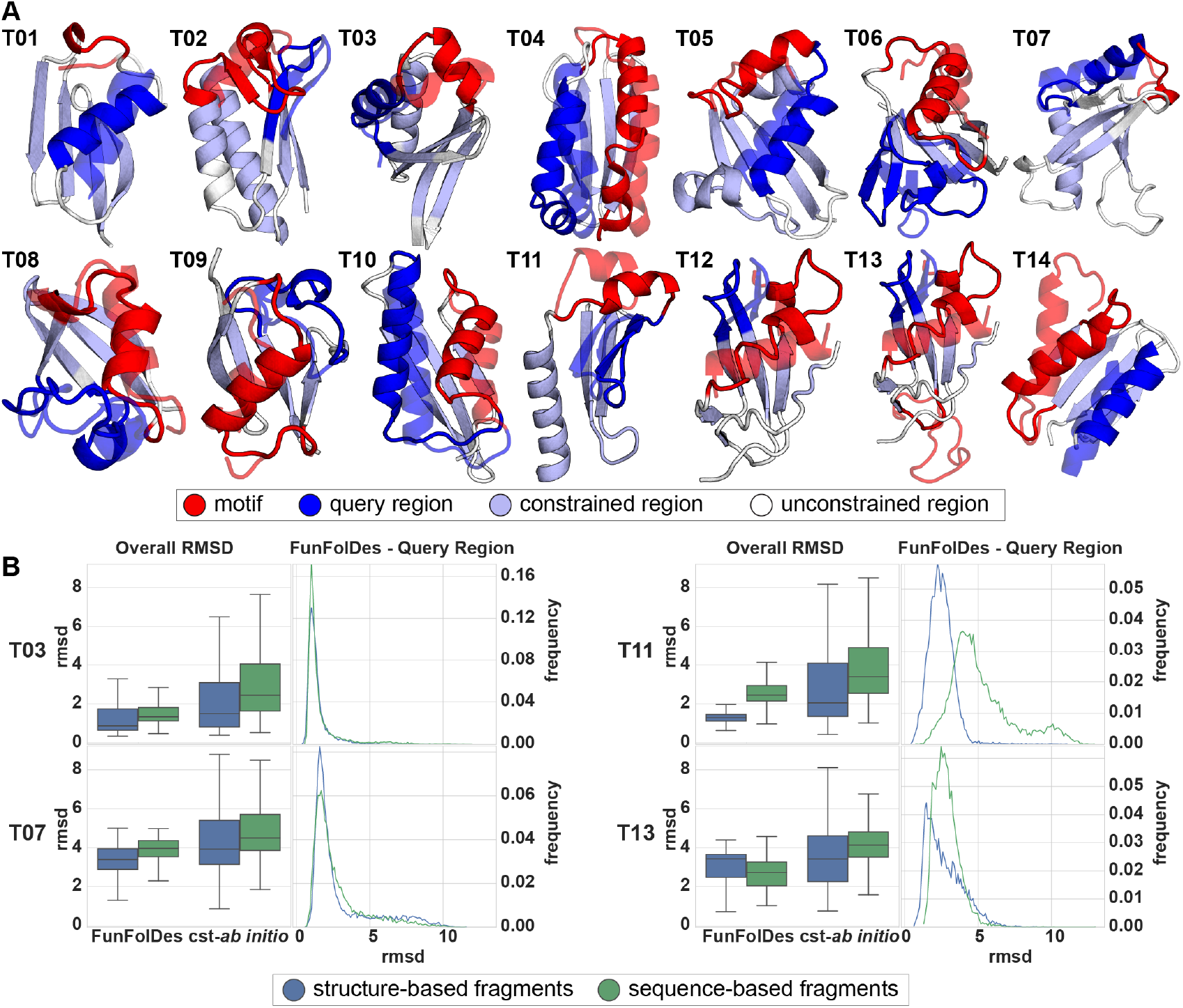
Benchmark test set to evaluate FunFolDes structural sampling. A) Structural representation of the 14 targets used in the benchmark. Each target highlights the motif (red), the query region (blue), and the positions from which distance constraints were generated (light blue). Conformations of the motif and query regions, as found in the template structures, appear superimposed in a semi-transparent depiction. B) Full structure RMSD (Overall RMSD) and local RMSD for the query region (FunFolDes - Query Region) for four targets (full dataset presented in Supplementary Figure S1). Overall RMSD compares results for the two simulation modes (FunFolDes Vs. constrained-*ab initio (cst-ab initio)*) and the two fragment generation methods (structure-(blue) Vs. sequence-based fragments(green)). FunFolDes more frequently samples RMSDs closer to the conformation of the target structure. Generally, structure-based fragment also contribute to lower mean overall RMSDs. The FunFolDes - Query Region RMSD distributions show that the two fragments sets do not have a major importance in the structural recovery of the query region.

Briefly, for the benchmark we selected proteins with less than 100 residues, where each benchmark test case is composed of two proteins of the same CATH domain cluster. One of the proteins is dubbed template, and serves as a structural representative of the CATH domain. The second protein, dubbed target, contains structural insertions or deletions (motif), to which a structural change in a different segment of the same structure could be attributed (query region). The motif and query regions for all the targets are shown in **Figure 2A** and quantified in terms of the percentage of overall secondary structure in **Figure 2 - Supplementary Figure 1A**. To a great extent, these structural changes due to natural sequence insertions and deletions are analogous to those occurring in the design scenarios for which FunFolDes was conceived.

Using FunFolDes, we folded and designed the target proteins while maintaining the motif segment structurally fixed, mimicking a structural motif insertion. Distance constraints between residues were extracted from the template in the regions of shared structural elements of the template and the target, and were used to guide the folding simulations.

To check whether FunFolDes enhances sequence and structural sampling, we compared the simulations to constrained *ab initio (cst-ab initio) simulations*(Bowers et al., 2000). These simulations were performed using the same sets of constraints but without the motif region as a static segment.

As Rosetta conformational sampling is highly dependent upon the fragment set provided, in this benchmark we also tested the influence of structure- and sequence-based fragments. The performance of the two protocols was broadly analyzed by the global and local recovery of both structure and sequence.

Structural recovery was assessed through two main metrics: (a) global RMSD of the full decoys against the target and (b) local RMSD of the query region. When evaluating the distributions for global RMSD in the designed ensembles, FunFolDes outperformed cst-*ab initio* by consistently producing populations of decoys with lower mean (RMSDs mostly found below 5 Å), a result observed in all 14 targets (**Figure 2B, Figure 2 - Supplementary Figure 1B**). This result is especially reassuring considering that FFL simulations contain more structural information of the target topology than the cst-ab *initio* simulations.

**Figure 2 - Supplementary Figure 1.**
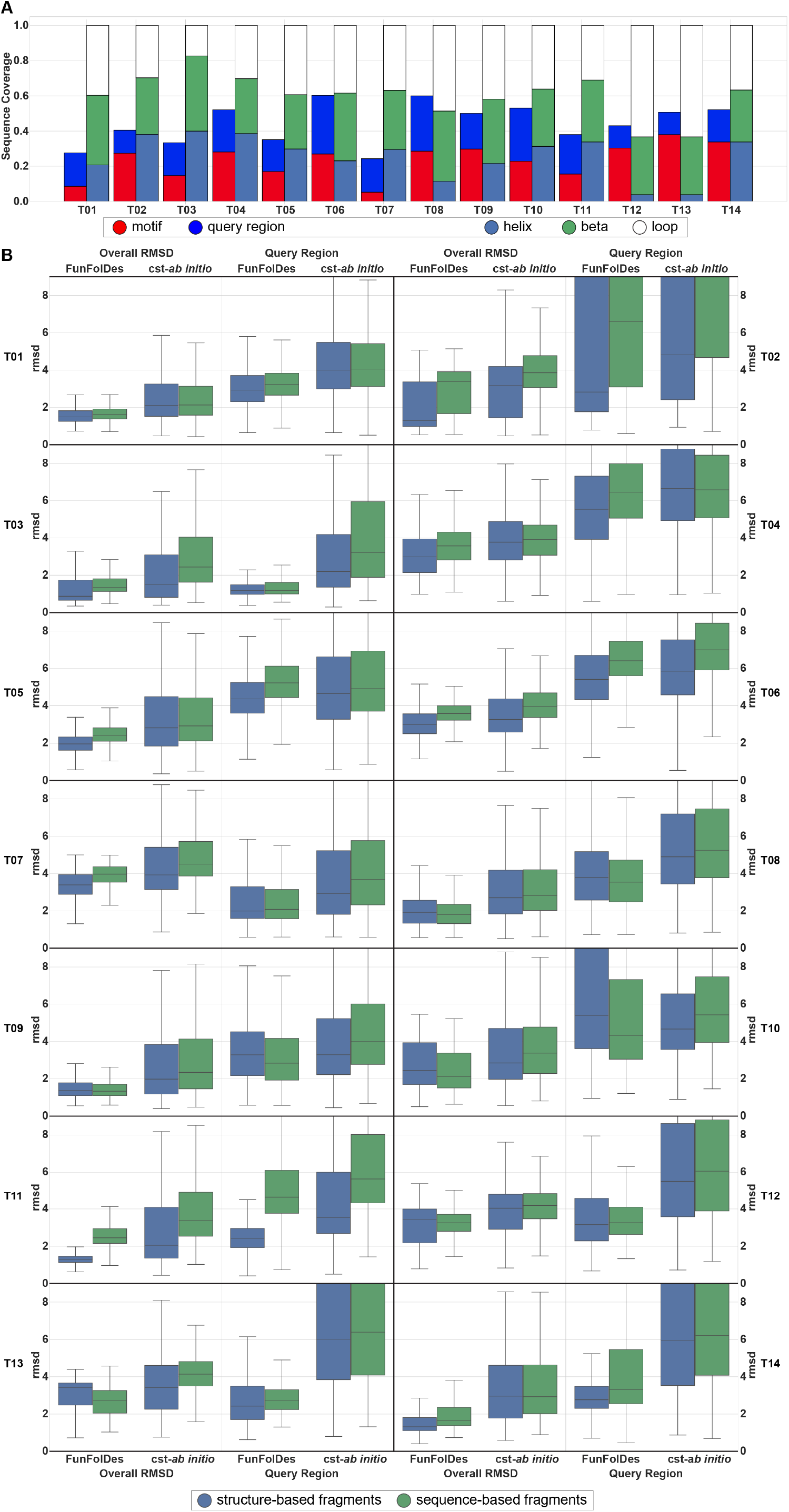
Structural composition and overall results of the benchmark targets. A) Percentage of secondary structure type, motif and query region in the overall structures. B) Full structure RMSD (Overall RMSD) and local RMSD for the query region (Query Region) between the decoy populations and their respective targets. FunFolDes tends to outperform *cst-ab initio* in all scenarios and the structure-based fragments yield decoy population with lower mean RMSDs, albeit with small differences relative to the sequence-based fragments.

Retrieval of the local RMSDs of the query unconstrained regions presented mixed results across the benchmark set. In 13 targets, FunFolDes outperforms cst-ab *initio*, showing lower mean RMSDs in the decoy population.

When comparing fragment sets (structure- vs sequence-based), both achieved similar mean RMSDs in the decoy populations; nonetheless, the structure-based fragments more often reach the lowest RMSDs for overall and query RMSDs (**Figure 2B, Figure 2 - Supplementary Figure 1**). This is consistent with what would be expected of structural information content within each set of fragments. When paired with the technical simplicity of use, time-saving and enhanced sampling of the desired topology, the structure-based fragments are an added value for FunFolDes.

In addition to structural metrics, we also quantified sequence recovery in the decoy populations, both in terms of sequence identity as well as sequence similarity according to the BLOSUM62 matrix (Henikoff & Henikoff, 1992) (**Figure 3A**). In all targets, the sequence identities and similarities were higher for FunFolDes populations than for cst-*ab initio*, and in line with sequence recoveries presented for other design protocols (Murphy et al., 2012) (**Figure 3A**). This type of metrics has been shown to be highly dependent on the exact backbone conformation used as input (Kuhlman & Baker, 2000; Murphy et al., 2012). Given that FunFolDes is exploring larger conformational spaces, as a proxy for the quality of the sequences generated, we used the target’s Hidden Markov Models (HMM) (Eddy, 2011) and quantified how many of the designed sequences were identified as belonging to the target’s CATH superfamily according to its HMM definition (**Figure 3B**).

**Figure 3.**
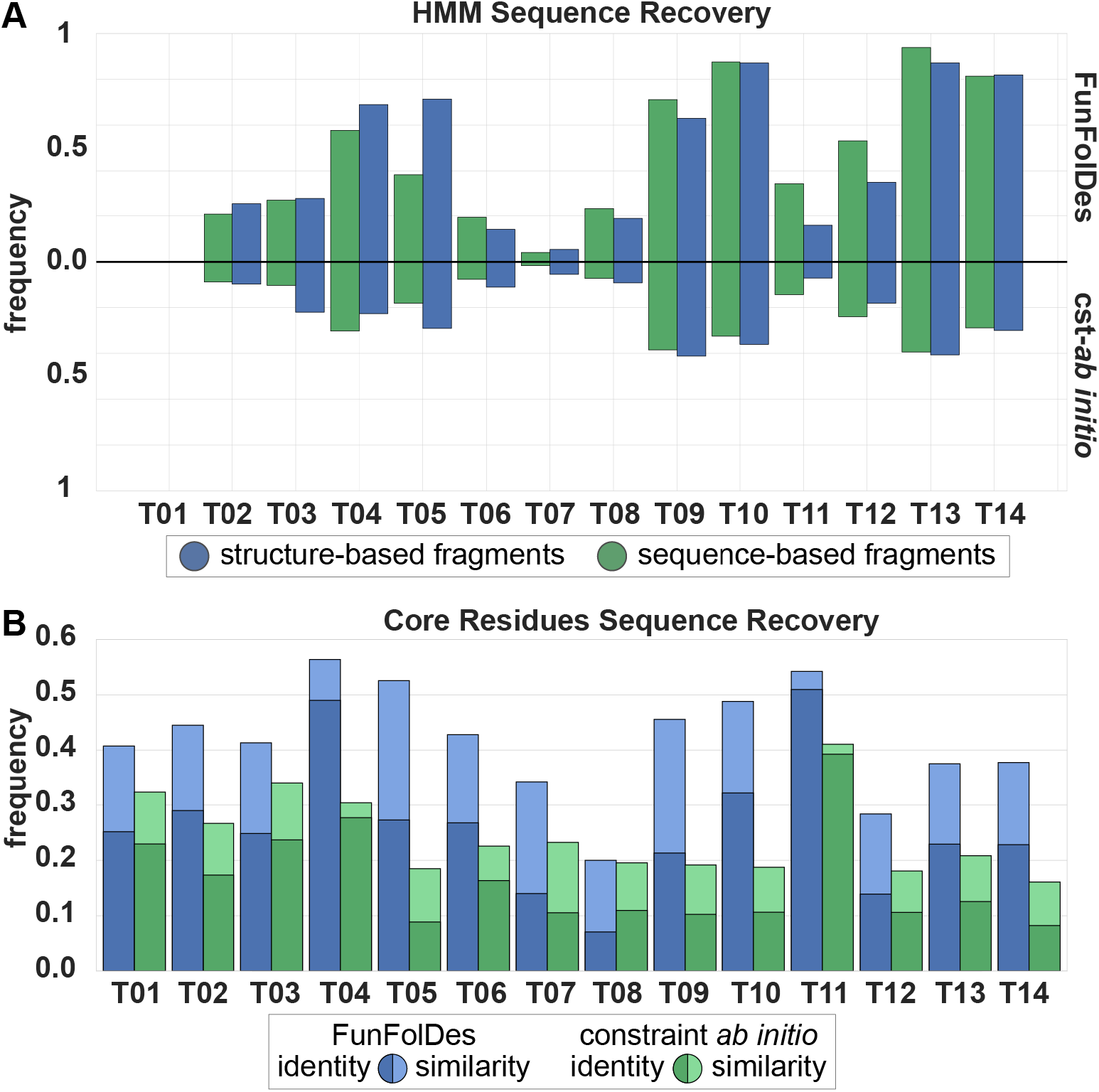
Assessment of FunFolDes’ sequence sampling quality of. A) HMM Sequence Recovery measures the percentage of decoys generated that can be assigned to the original HMM from the CATH superfamily that the target belongs to. FunFolDes consistently outperforms cst-*ab initio*, which is consistent with the same behavior observed in the structural recovery. B) Core Residues Sequence Recovery reveals the conservation of core residues between each design set and its target. Conservation is measured in terms of sequence identity and sequence similarity (as assigned through BLOSUM62). Also according to this metric FunFolDes outperforms cst-ab *initio* in every instance, reaching, for some populations, levels of conservation similar to those found in more restrained flexible-backbone designs.

HMM recovery was computed as the percentage of decoys with an E-value under 10 and covering more than 50% of the full decoy sequence. FunFolDes decoy populations systematically outperformed those from cst-ab *initio* (**Figure 3B**). The performance of the two fragment sets shows no significant differences. Core sequence identity and similarity was assessed over the structure-based fragment set.

In summary, the results of this benchmark highlight the usability of FunFolDes to generate close-to-native scaffold proteins to stabilize inserted structural motifs. FunFolDes aims to refit protein scaffolds towards the requirements of a functional motif. In this perspective, it is critical to explore, within certain topological boundaries, structural variations around the original template. This benchmark points to several variables in the protocol that resulted in enhanced structural and sequence sampling.

### Target-biased folding and design of protein binders

The computational design of proteins that can bind with high affinity and specificity to targets of interest remains a largely unsolved problem (Schreiber & Fleishman, 2013). Within FunFolDes’ conceptual approach of coupling folding with sequence design, we sought to add the structure of the binding target (**Figure 1**) to attempt to bias sampling towards functional structural and sequence spaces.

Previously, we used FFL to design a new binder (BINDI) to BHRF1 (**Figure 4A**), an Epstein-Barr virus protein with anti-apoptotic properties directly linked to the tumorigenic activity of EBV (Procko et al., 2014). FFL was used to generate the initial designs that bound to BHRF1 with a dissociation constant (*K*_D_) of 5860 nM, which were then affinity matured (*K*_D_ = 220±50 pM) and showed improved bacterial expression. BINDI was designed in the absence of the target and then docked to BHRF1 through the known interaction motif. The BHRF1-compatible models were further designed to ensure structural compatibility and improve affinity. A striking observation from the overall approach was that the FFL stage was highly inefficient, generating a large fraction of backbone conformations incompatible with the binding mode of the complex.

**Figure 4.**
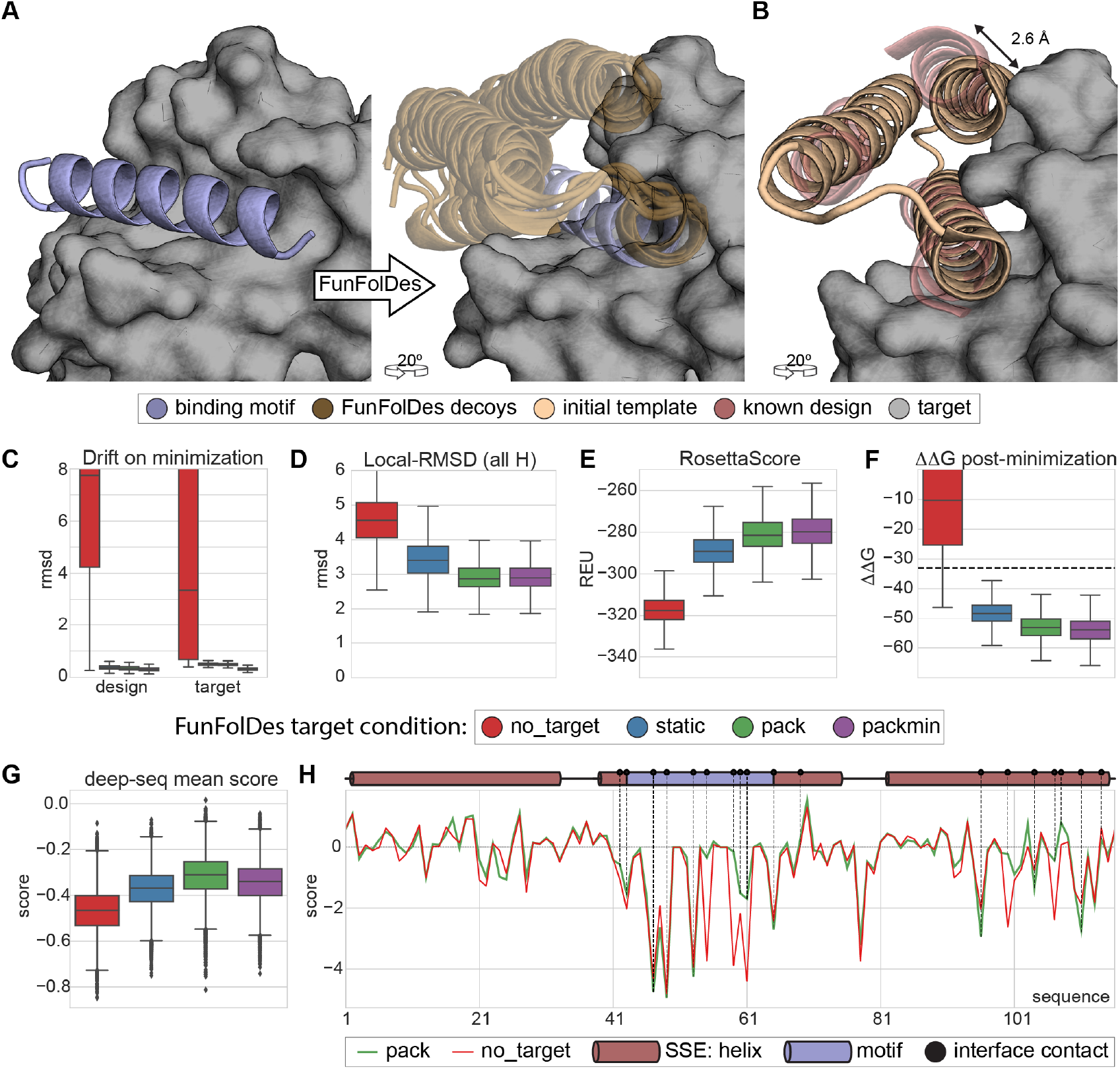
Target-biased design of a protein binder and assessment of performance based on saturation mutagenesis data. A) Depiction of the initial design task, a single-segment binding motif (BIM-BH3), shown in purple cartoon representation, with its target (BHRF1), shown in gray surface is used by FunFolDes to generate an ensemble of designs compatible with the binding mode. B) Conformational difference between the initial template (PDB ID: 3LHP) and the previously designed binder (BINDI shown in pink cartoon representation), helix 3 requires a subtle but necessary shift (2.6 Å) to avoid steric clashes with the target. C-G) Scoring metrics for each design population according to the simulation mode: no_target - FunFolDes was used without the target protein; static - target present no flexibility allowed; pack - target allowed to repack the side-chains; packmin – side-chain repacking plus minimization and backbone minimization were allowed for the target. The target flexibility was allowed during the relax-design cycle of FunFolDes. C) Structural drift observed for design and target binder measured as the RMSD difference of each structure between pre- and post-minimization conformations. D) Structural recovery of the conformation observed in the BINDI-BHRF1 assess over the 3 helical segments of the bundle. E) Rosetta energy for the design populations generated with different simulation modes. F) Interaction energy (ΔΔG) between the designs and the target. G) Deep-sequence score distribution for each design population, computed as the mean score of each sequence after applying a position score matrix based on the deep-sequence data. The pack population slightly outperforms the other simulation modes. H) Per-residue scoring comparison of the no_target and the pack populations according to the deep-sequence data. Although the behavior is overall similar, pack outperforms no_target in multiple positions, several of which are highlighted(black dots) as interfacial contacts or second shell residues close to the bind site which were allowed to be designed throughout the simulations.

To test whether the presence of the target could improve structural and sequence sampling, we leveraged the structural and sequence information available for the BINDI-BHRF1 and benchmarked FunFolDes for this design problem.

As described by Procko and colleagues, when comparing the topological template provided to FFL and the BINDI crystal structure, the last helix of the bundle (helix 3) was shifted relative to the template to ensure structural compatibility between BINDI and BHRF1 (**Figure 4B**). We used this case study to assess the capabilities of FunFolDes to sample closer conformations to those observed in the BINDI-BHRF1 crystal structure. In addition, we compared the saturation mutagenesis data generated for BINDI (Procko et al., 2014) to evaluate the sequence space sampled by FunFolDes.

A detailed description of this benchmark can be found in the Materials and Methods section. Briefly, we performed four different FunFolDes simulations: I) binding target absent (no_target); II) binding target present with no conformational freedom (static); III) binding target present with side-chain repacking (pack); IV) binding target present with side-chain repacking plus minimization and backbone minimization (packmin). After the FunFolDes simulations, the no_target set was docked to BHRF1 through the binding motif and the remaining three simulations produced complexed structures. All the complexes were globally minimized (both design and target) to assess the conformational and energy changes as a proxy of the structural compatibility of the designed binders.

Simulations performed with the target absent (no_target) very rarely produce conformations compatible with the target (<10% of the total generated designs) (**Figure 4 - Supplementary Figure 1A**). We observed an improvement on the fraction of decoys compatible with the binding target (>60%) after global minimization (**Figure 4 - Supplementary Figure 1A**). However, this was at the cost of considerable structural drifts for both binder (mean RMSD 3.3 Å) and target (mean RMSD 7.7 Å) (**Figure 4C**). These structural drifts are a reflection of the energy optimization requirements by the relaxation algorithms but tat are deemed biologically irrelevant due to the profound structural reconfigurations. In contrast, simulations performed in the presence of the target clearly biased the sampling to more productive conformational spaces. RMSD drifts upon minimization were less than 1 Å for both designs and target (**Figure 4C**).

**Figure 4 - Supplementary Figure 1.**
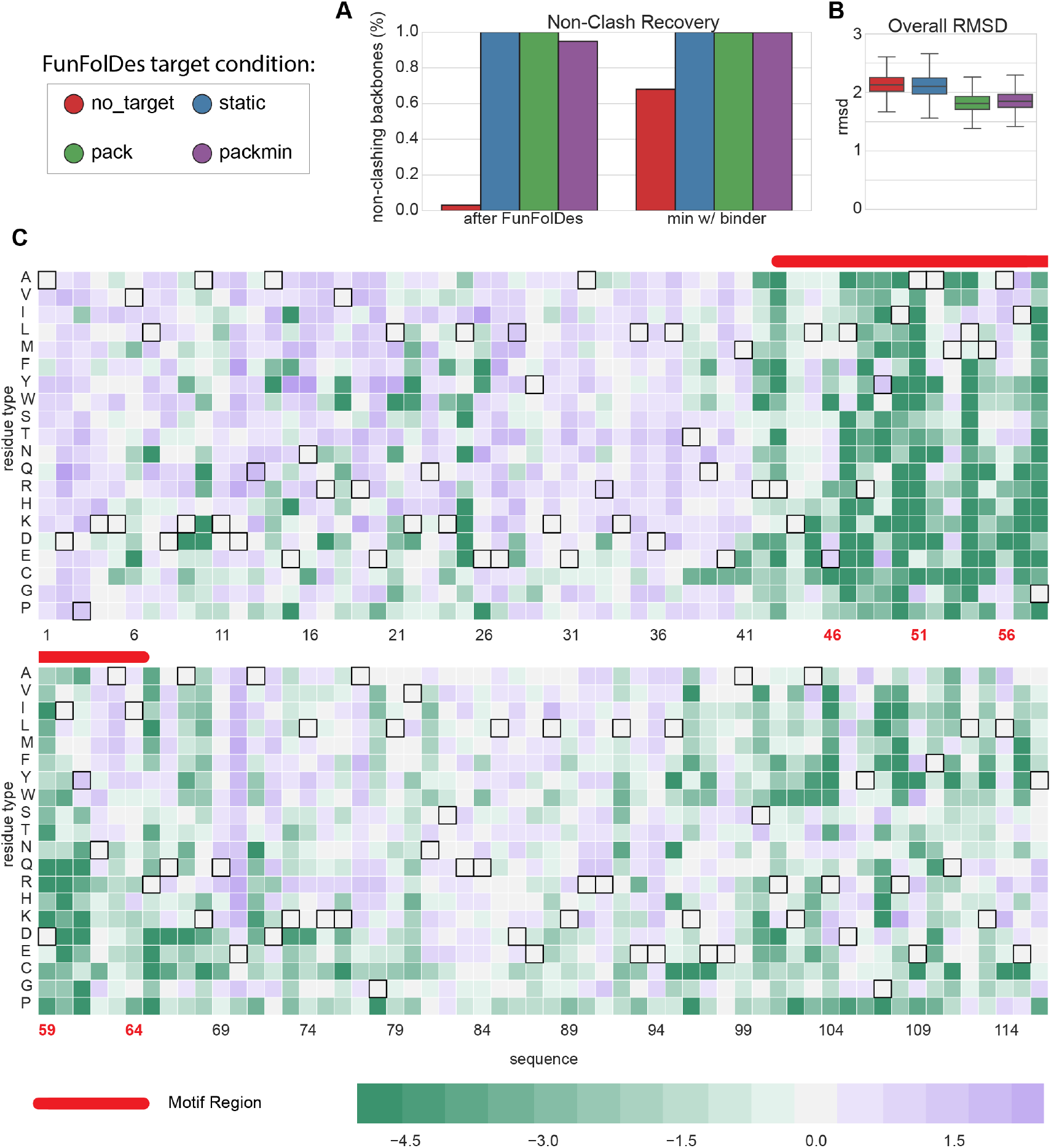
Target-biased folding and design: structural features od the modeled designs and saturation mutagenesis data used for sequence recovery benchmark. A) Quantification of the percentage of decoys compatible with a design-target binding conformation for the different simulation modes. The simulations performed without the target yield a very low percentage of binding compatible conformations. After minimization, this percentage increases with significant structural drifts. B) The initial template is a 3-helix bundle structure, the slight shift needed to adopt a binding-compatible conformation produces only a small global RMSD. C) Graphical representation of the deep-sequencing data as a position-specific score matrix. Black borders highlight the native BINDI residue type for each position. Mutations for which no data was obtained, likely reflect that these protein variants were unable to fold an display at the surface of yeast and were assigned the lowest score of −5.

Given that global alignments of the designs do not emphasize the local differences and the helical arrangement (**Figure 4 - Supplementary Figure 1**), to analyze structural regions of particular interest, we aligned all the designs on the conserved binding motif (**Figure 4A**) and measured the RMSD over the three helices that compose the fold. The two key regions were helices 1 and 3, which are in direct contact with the target.

According to this metric, FunFolDes simulations in the presence of the target sampled mean RMSD of 3 Å with the BINDI structure as reference (**Figure 4D**), and the closest designs were at approximately 2 Å. On the other hand, simulations in the absence of the target showed a mean RMSD of 4.5 Å, and the best designs around 2.5 Å.

While we acknowledge that these structural differences are modest, the data in this benchmark suggest that these differences can be important in the sampling of conformations and sequences competent for binding.

In addition to structural sampling, we also analyzed Rosetta Energies for the different simulations. We observed noticeable differences in the overall energy of the designed binders; in the absence of the binding target, the designs have an mean energy of approximately −320 Rosetta Energy Units (REUs), while the designs generated in the presence of the binding target showed an mean between −280 and −290 REUs (**Figure 4E**). This difference is significant, particularly for a protein of such small size (116 residues). Likewise, we also observe considerable differences in terms of the binding energy (ΔΔG) for the designs folded in the absence or in the presence of the binding target, corresponding to mean ΔΔGs of −10 and −50 REUs, respectively (**Figure 4E**).

The energy metrics provide interesting insights regarding the design of functional proteins. Although the sequence and structure optimization for the designs in the absence of the target reaches lower energies, these designs are structurally incompatible with the binding target and, even after refinement, their functional potential (as assessed by the ΔΔG) is not nearly as favorable as those performed in the presence of the binding target (**Figure 4F**). These data suggest that, in many cases, to optimize function it may be necessary to sacrifice the overall computed energy of the protein which is often connected to the experimental thermodynamic stability of the protein. Although stability is an essential requirement for all functional proteins (Chevalier et al., 2017; Tokuriki, Stricher, Serrano, & Tawfik, 2008), it may be necessary to design proteins that are, *in silico*, less energetically favorable to ensure that the target functional requirements can be accommodated. This observation provides a compelling argument to perform biased simulations in the presence of the binding target, which may broadly be defined as a “functional constraint”.

To evaluate sequence sampling quality, we compared the computationally designed sequences to a saturation mutagenesis dataset available for BINDI (Procko et al., 2014). Briefly, this dataset was obtained by screening a saturation mutagenesis library for binding interactions in a yeast-display setup coupled to a deep sequencing readout. The impact on the binding affinity of each mutation was assessed based on the relative frequencies of the mutants. Data from this experiment were transformed into a positional scoring matrix (**Figure 4 - Supplementary Figure 1C**). Point mutations that showed a beneficial effect on the binding affinity to BHRF1 have a positive score, deleterious mutants a negative score, and neutral score 0. Such a scoring scheme, will yield a score of 0 for the BINDI sequence.

When scoring the designs generated by the four different simulations, designs performed in the presence of the binding target obtain higher mean scores as compared to the no_target designs (**Figure 4G**). The pack simulation, where the binding target is simply repacked, is the best performer with the highest distribution mean, having one design that scores better than the BINDI sequence. Furthermore, it is important to highlight that in some key positions at the protein-protein interface, the pack designs clearly outperformed those generated by the no_target simulation, when quantified in terms of a per-position score (**Figure 4H**); meaning that across the design population, amino-acids that can be conducive to productive binding interactions were sampled more often in the presence of the binding target. This sequence sampling benchmark provides an example of the benefits of using a “functional constraint” (binding target) to improve the quality of the sequences obtained by computational design.

Overall, the BINDI benchmark provides important insights regarding the best computational protocol within FunFolDes that can be utilized to improve the outcome of design simulations in terms of frequency of functional proteins.

### Repurposing a naturally occurring fold for a new function

At its conception, FFL was primarily envisioned to aid the design of function into proteins. To further test FunFolDes’ design capabilities, we sought to transplant a contiguous viral epitope that can be recognized by a monoclonal antibody with high affinity (**Figure 5A**). The success of the designs was assessed by their folding, thermal stability, and more importantly, binding affinities to the epitope-specific antibody as the functional readout.

**Figure 5.**
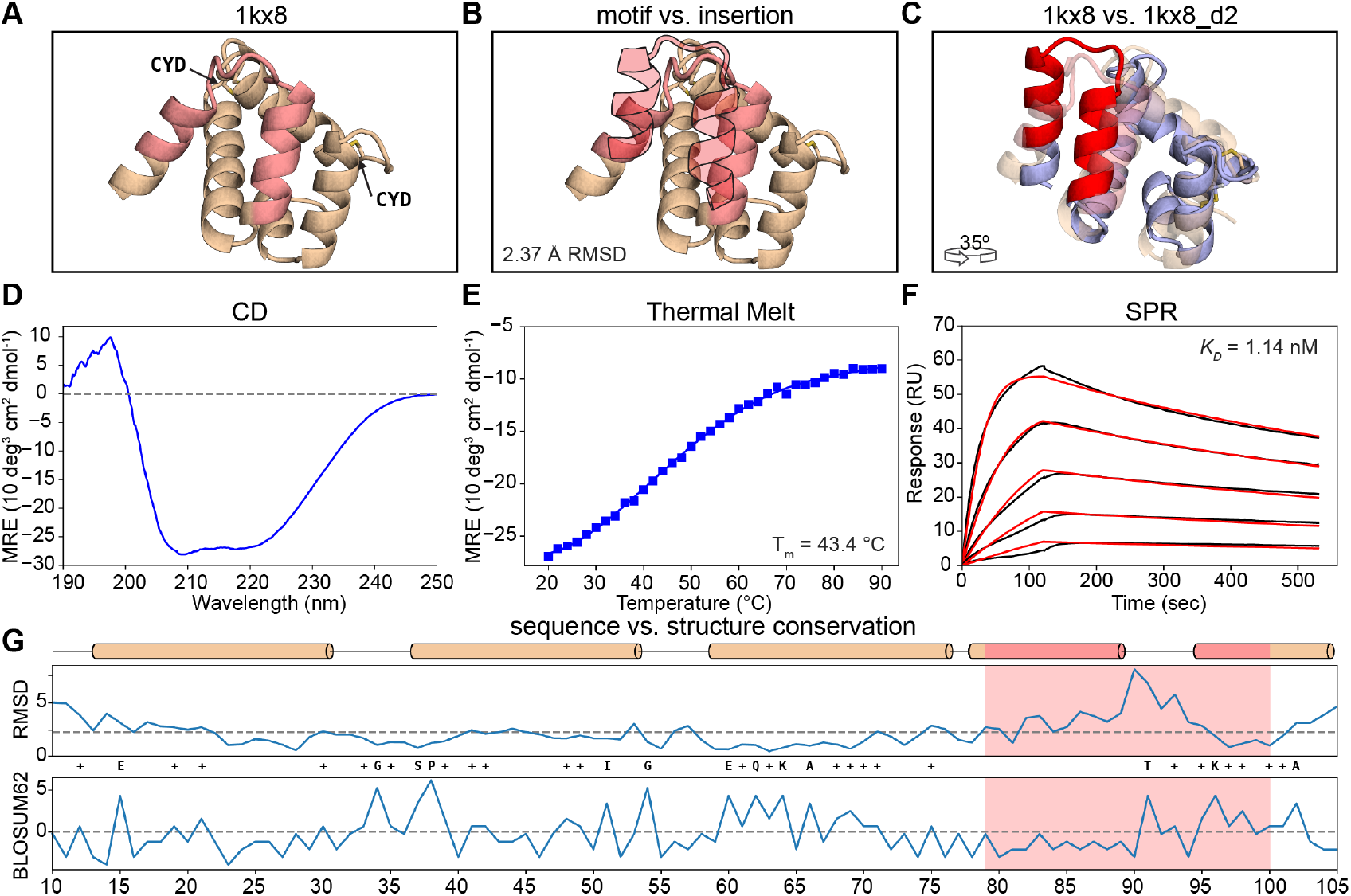
Functional design of a distant structural template. A) Structural representation of 1kx8. The insertion region is colored in light red and the two disulfide bonds are labeled (CYD). B) Structural comparison between the insertion region of 1kx8 and the mota epitope (light red-filled silhouette). Local RMSDs between the two segments reach 2.37 Å. C) Superimposition between 1kx8_d2 design model (blue with red motif) and the 1kx8 template (wheat and light red insertion site). Multiple conformational shifts are required throughout the structure to accommodate the site II epitope. D) CD spectrum of 1kx8_d2 showing a typical alpha-helical pattern with the ellipticity minimums at 208 nm and 220 nm. E) 1kx8_d2 shows a melting temperature (T_m_) of 43.4°C. F) Binding affinity determined by SPR. 1kx8_d2 shows a Kd of 1.14 nM. Experimental sensorgrams are shown in black and the fitted curves in red. G) Per-position evaluation of structural (top) and sequence (bottom) divergence between the design model 1kx8_d2 and the starting template 1kx8. The largest structural differences are observed in the region where the site II epitope was inserted, the overall difference of the two structures is 2.25 Å (dashed line). Sequence divergence is evaluated by applying the BLOSUM62 score matrix to the sequences, yielding a total of 13.5% identity and 38.5% similarity mostly in the structured regions. The epitope region is colored in light red. Identical positions between the 1kx8_d2 and 1kx8 are labeled with the residue one letter code while positively scored changes according to BLOSUM62 are labeled with a + symbol.

Specifically, we used as functional motif the RSVF site II epitope (PDB ID: 3IXT (McLellan, Chen, Kim, et al., 2010)), a helix-loop-helix motif recognized by the antibody motavizumab (mota). Previously, we have designed proteins with this same epitope (Correia et al., 2014); however, we started from a structural template with a similar conformation to that of the epitope, the RMSD between the epitope and the scaffold segment was approximately 1 Å when measured over the helical positions. Here, we sought to challenge FunFolDes by using a distant structural template where the local RMSDs of the epitope structure and the segment onto which the epitope was transplanted would be over 2 Å. We used master (Zhou & Grigoryan, 2015) to perform a structural search of the site II epitope over a subset of structures in the PDB. After filtering the results by scaffold size (50-100 residues) and steric clashes using the structure of the epitope in complex with mota, we selected as template scaffold the structure of the A6 protein of the Antennal Chemosensory system from the moth *Mamestra brassicae* (PDB ID: 1KX8 (Lartigue et al., 2002))(**Figure 5A**). The backbone RMSD between the conformation of the epitope and the insertion region in 1kx8 is 2.37 Å (**Figure 5B**).

In terms of biological function, 1kx8 is involved in chemical communication and perception(Lartigue et al., 2002). Biochemically, it has been shown to bind to fatty-acid molecules with hydrophobic alkyl chains composed of 12-18 carbons. Two prominent features are noticeable in the structure of 1kx8: two disulfide bonds (**Figure 5A**) and a considerable void volume in the protein core, deemed to be the binding site for fatty acid molecules. These features emphasize that the initial design template is likely not a very stable protein.

In the design process we performed two stages of FunFolDes simulations; first an exploratory stage to select properly folded designs with the functional motif inserted (**Figure 5C**) that were fed into a second round of simulations, which sampled much more extensively within the structural proximity of the 1^st^ generation template. For each stage, we generated 12’500 designs, eventually selecting seven for initial experimental characterization according to several structural features of the computational models, namely: Rosetta Energy, packing score, and buried unsatisfied hydrogen bonds (**Figure 5 - Supplementary Figure 1**). A detailed description of the process can be found in the Materials and Methods.

**Figure 5 - Supplementary Figure 1.**
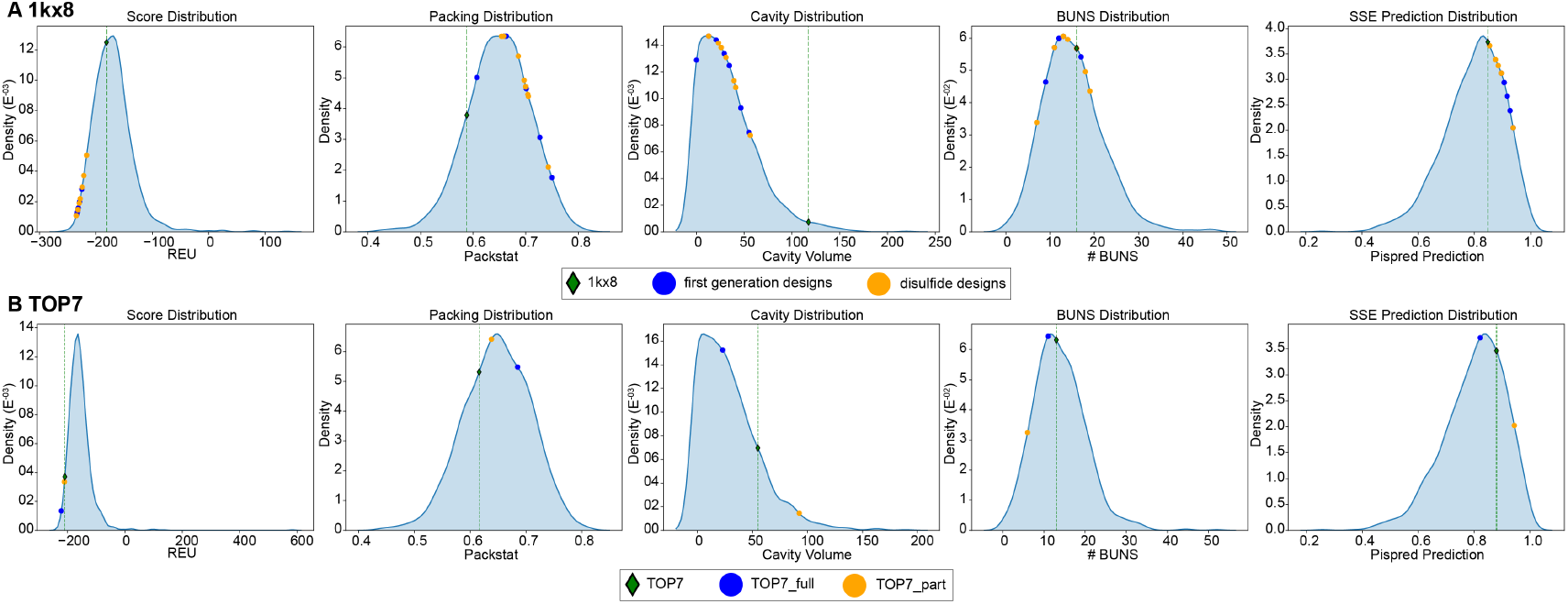
Structural and sequence evaluation of the computational designs. Assessment of structural and sequence features: Rosetta Energy, packing score (packstat) (Alford et al., 2017), cavity volume, Buried UNSatisfied polar atoms and secondary structure prediction (PSIPRED) for the template and the computational designs. Each template (green diamond) and design (yellow and blue circles) are compared against a set of non-redundant minimized structures of similar size (± 15 residues). A) Due to its natural function, 1kx8 presents of a large cavity to bind its hydrophobic ligands. As such, the structure presents generally low scores as compared to computationally designed proteins. B) Distributions of the structural and sequence features of natural proteins and the TOP7 series of designs.

We characterized experimentally the computationally designed; those expressed in bacteria at good yields were further characterized using size exclusion chromatography coupled to a multi-angle light scatter (SEC-MALS) to determine the solution oligomerization state. To assess their folding and thermal stability(Tm) we used Circular Dichroism (CD) spectroscopy, and finally to assess their functional properties we used surface plasmon resonance (SPR) to determine binding dissociation constants (*K_D_*s) to the mota antibody. We started by expressing seven sequences from our first round of computational design; out of these seven, six designs were purified and characterized further. While the majority of the designs were monomers in solution and showed CD spectra typical of helical proteins, in terms of stability we obtained both designs that were not very stable nor did they exhibit cooperative unfolding (1kx8_02) and also designs that were very stable and did not fully unfold under high temperatures (1kx8_07) (**Figure 5 - Supplementary Figure 2**).

**Figure 5 - Supplementary Figure 2.**
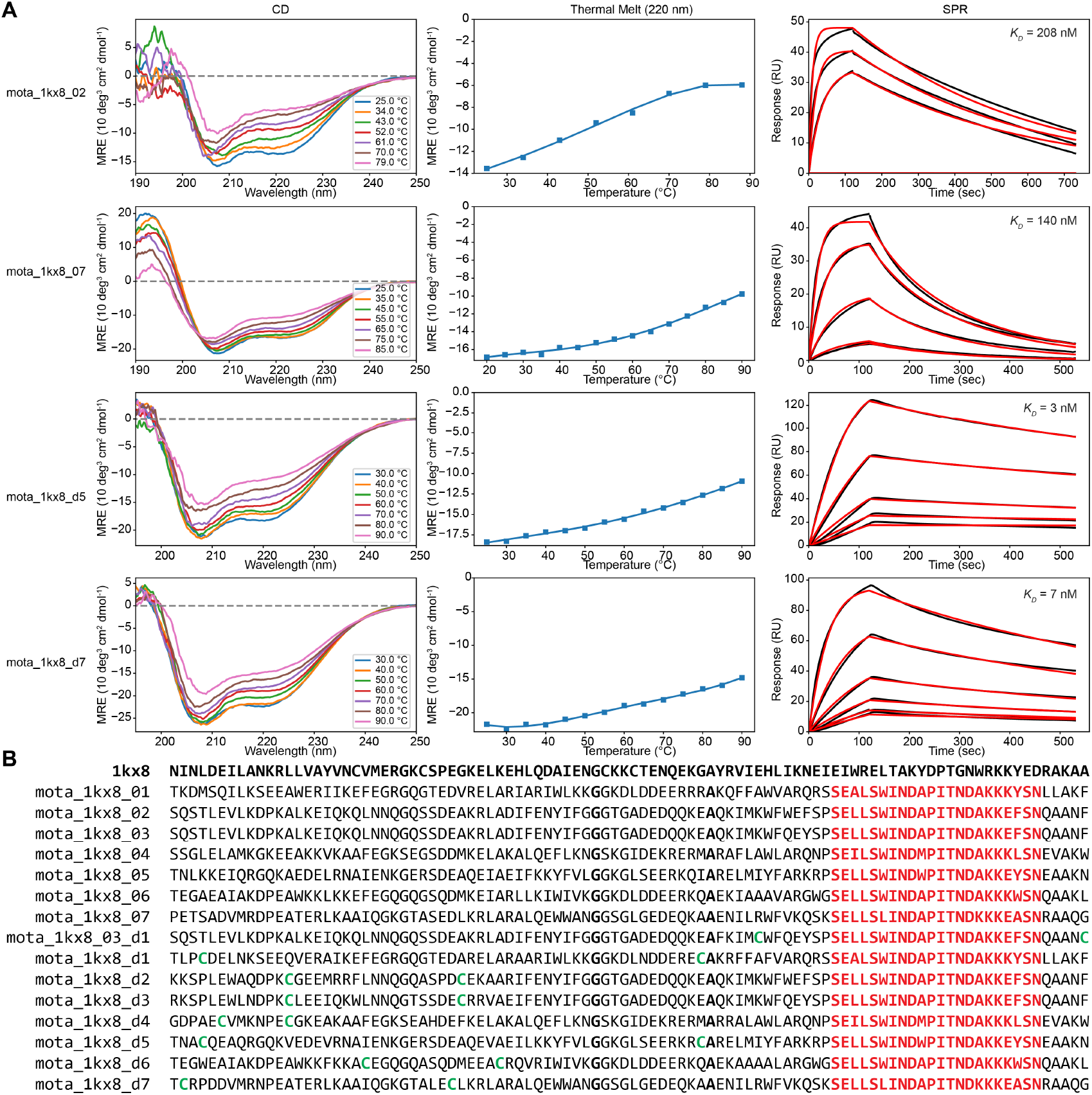
Examples of experimental characterization performed for other variants on the 1kx8 design series. A) CD wavelength spectra (left column), thermal denaturations (middle column) and SPR binding assays with the mota antibody (right column) were performed. B) Global sequence alignment of the wild-type protein 1kx8 and the computationally designed sequences. Red positions highlight the site II epitope insertion. Green positions highlight the cysteines performing the disulfide bridges. The two positions that consistently kept the original residue type of 1kx8 are highlighted in bold.

The determined binding affinities to mota ranged from 34 to 208 nM, which was an encouraging result. Nevertheless, comparing this affinity range to those of the peptide epitope (*K*_D_ = 20 nM) and other designs with the same site grafted that were published previously (*K*_D_ = 20 pM) (Correia et al., 2014), there was room for improvement. Therefore, we generated a second round of designs to attempt to improve stability and binding affinities. Driven by the observation that the native fold has two disulfides bonds, our next set of designs included engineered disulfide bonds.

In the second round, we tested eight designed variants with different disulfide bonds and, if necessary, additional mutations to accommodate them. All eight designs were soluble after purification and two were monomeric: 1kx8_d2 and 1kx8_3_d1, which also showed CD spectra typical of helical proteins (**Figure 5D**) with melting temperatures (T_m_s) of 43 and 48°C (**Figure 5E**), respectively. Remarkably, 1kx8_d2 showed a *K*_D_ of 1.14 nM (**Figure 5F**), an improvement of approximately 30-fold compared to the best variants of the first round. 1kx8_d2 binds to the mota with approximately 20-fold higher affinity than the peptide-epitope (*K*_D_ ≈ 20 nM), and 50 fold lower compared to previously designed synthetic scaffolds (*K*_D_ = 20 pM) (Correia et al., 2014). This difference in binding is likely reflective of how challenging it can be to accomplish the repurposing of protein structures with distant structural similarity.

Post-design analyses were performed to compare the sequence and structure of the best design model with the initial template. **Figure 5G**, shows a per-residue RMSD measurement upon a global alignment of the 1kx8 structure with the designed model. The global RMSD between the two structures is 2.25 Å. Much of the structural variability arises from the inserted motif, while the surrounding segments adopt a configuration similar to the original template scaffold. The sequence identity of 1kx8_d2 as compared to the native protein is approximately 13%. The sequence conservation per-position (**Figure 5G**) was evaluated through the BLOSUM62 matrix, where positive scores are attributed if the original residue is not mutated or if the substitution is deemed favorable according the scoring matrix, and negative if unfavorable. Overall, 38.5% of the residues in 1kx8_d2 scored positively, and 61.4% of the residues had a score equal to or lower than 0. This is particularly interesting in the perspective that multiple mutations deemed unfavorable according the statistics condensed in the BLOSUM62 matrix are still able to yield well folded and, in this case, functional proteins.

The successful design of this protein is a relevant demonstration of both the broad usability of the FFL algorithm and of the overall strategy of designing functional proteins by coupling the folding and design process to incorporate functional motifs in unrelated protein folds. In a subsequent design challenge, we sought to functionalize a *de novo* design fold, which unlike natural proteins, did not evolve under any sort of functional pressure.

### Functionalization of a functionless fold

Advances in computational design methodologies have achieved remarkable results in the design of *de novo* protein sequences and structures (Hill et al., 2000; Koga et al., 2012; Marcos et al., 2017). However, the majority of the designed proteins are “functionless” and were designed to test the performance of computational algorithms in predicting structural accuracy. Here, we sought to use one of the hallmark proteins of *de novo* design efforts - TOP7 (Kuhlman et al., 2003) (**Figure 6A**) - and functionalize it using FunFolDes. To do so, we leveraged several of the newly implemented features in FunFolDes. The functional site selected to insert into TOP7 was another viral epitope from RSVF, commonly referred to as site IV, which is recognized by the 101F antibody (McLellan, Chen, Chang, et al., 2010). When bound to the 101F antibody, site IV adopts a β-strand-like conformation (**Figure 6B**), which in terms of secondary structure content is compatible with one of the edge strands of the TOP7 topology (**Figure 6C**). Despite the secondary structure similarity, the RMSD of the site IV backbone in comparison with that of TOP7 is 2.1 Å, and upon alignment of the antibody in this particular orientation, clashes arise between TOP7’s helix 1 and the antibody interface. Therefore, this design challenge is yet another prototypical application for FunFolDes. In this design challenge we followed two distinct routes: I) a conservative approach where we fixed the amino-acid identities of roughly half of the core of TOP7 and allowed mutations mostly on the contacting shell of the epitope insertion site; and II) a sequence unconstrained design where all the positions of the scaffold were allowed to mutate. We attempted five designs for recombinant expression in *E. coli* and two (TOP7_full and TOP7_partial) were selected for further biochemical and biophysical characterization, one from each of the two design strategies mentioned above. According to SEC-MALS, both behaved as monomers in solution, with TOP7_partial being a less well-behaved protein with higher aggregation propensity. Both TOP7_full and TOP7_partial (Supplementary **Figure 5**) were folded according to CD measurements, with the TOP7_full showing a CD spectrum (**Figure 6D**) which very closely resembles that of the native TOP7 (Kuhlman et al., 2003). TOP7_full was subjected to thermal denaturation monitored by CD, where we observed that the newly designed protein is much less stable than the original TOP7 (**Figure 6E**). To quantify the functional component of TOP7_full, we determined the *K*_D_ of its interaction with 101F to be 24.2 nM (**Figure 6F**), which is within the range measured for the native viral protein RSVF (3.6 nM) (McLellan, Chen, Chang, et al., 2010). Importantly, the *K*_D_ for TOP7_full is 2400 fold higher than that of the peptide-epitope (58.4 μM) (McLellan, Chen, Chang, et al., 2010), suggesting that productive conformational stabilization and/or extra contacts to the rest of the protein were successfully designed.

**Figure 6.**
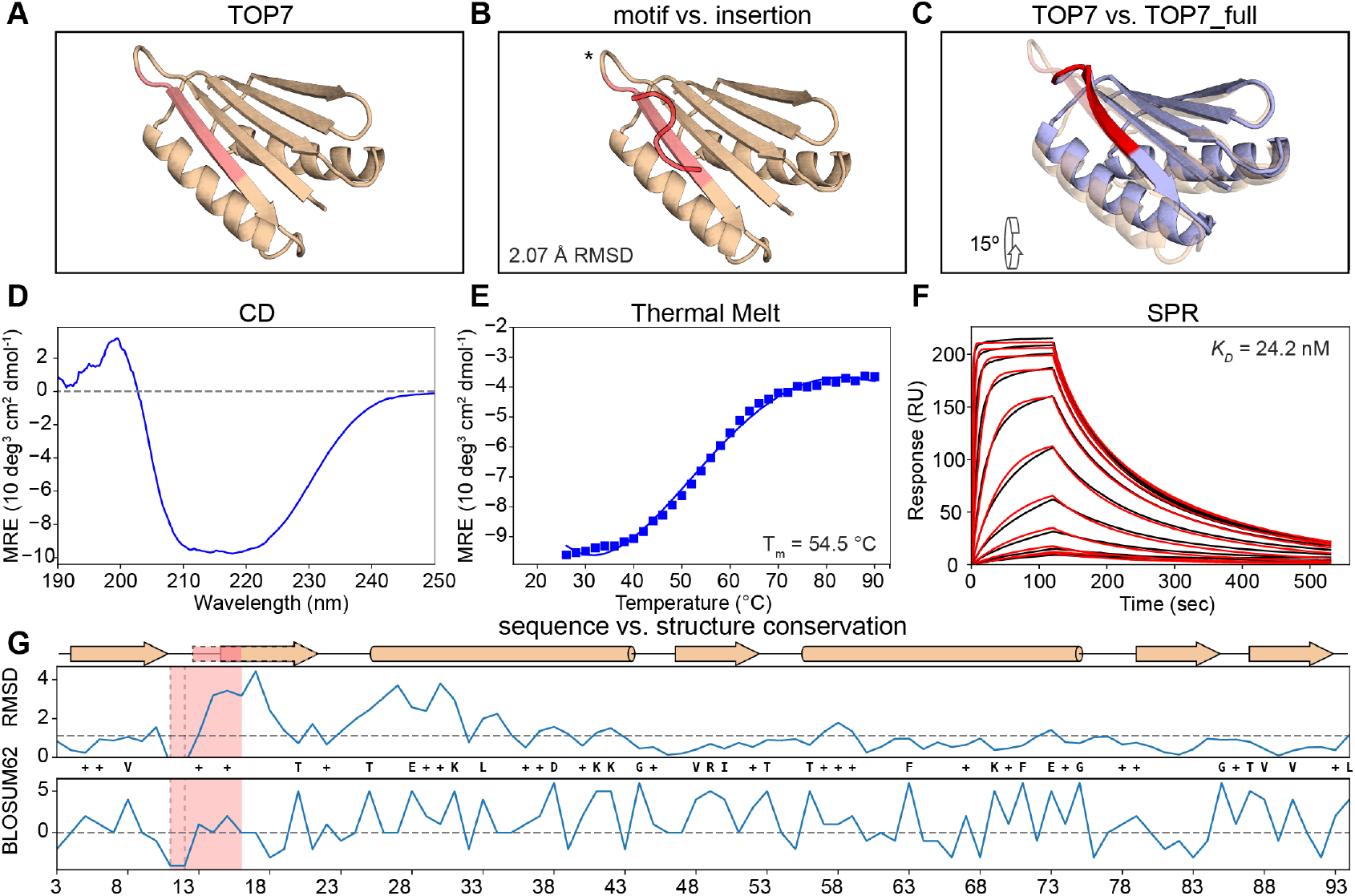
Functionalization of the functionless de novo fold TOP7. A) Structure of TOP7 with the insertion region highlighted in light red. B) Structural comparison between 101F and the insertion region of TOP7 reveals a 2.07 Å RMSD. C) TOP7_full model (in blue and red for the motif) superimposed over the TOP7 crystal structure. 101F’s insertion is structurally compensated mostly by the first pairing beta strand and a shift of the first alpha helix D) CD spectrum shows a broad ellipticity signal between 210 nm and 222 nm as a representative of mixed secondary structural propensities. E) The melting temperature (T_m_) for TOP7_full was 54.5 °C. F) Binding affinity determined by SPR. TOP7_full shows a *K*_D_ of 24.2 nM. Experimental sensorgrams are shown in black and the fitted curves in red. G) Per-position evaluation of structural (top) and sequence (bottom) divergence between the design model TOP7_full and the starting template TOP7. The largest structural differences are observed in the region downstream of the site IV epitope, the overall difference of the two structures is 1.5 Å (dashed line). Sequence divergence is evaluated by applying the BLOSUM62 score matrix to the sequences, yielding a total of 27.7% identity and 52.2% similarity. The epitope region is colored in light red. Identical positions between the TOP7_full and TOP7 are displayed as their residue types while positively scored changes according to BLOSUM62 are labeled with a + symbol.

Given the successful functionalization of TOP7, we sought to understand the levels of sequence and structural change (**Figure 6G**). Per-residue sequence recovery and structural similarity were evaluated for TOP7_full against TOP7 (**Figure 6G**). We compared the per-residue RMSD between the TOP7_full model and the crystal structure of TOP7, revealing that most conformational changes occur from the site IV insertion region and displacement of the neighboring alpha-helix, with the overall backbone RMSD between both structures being 1.5 Å. The connecting loop between the strand that holds the epitope and the adjacent strand was also shortened to obtain a tighter connection between the 2 strands (**Figure 6C**).

Remarkably, the sequence identity of the most aggressive design (TOP7_full) is only 28%, and using the BLOSUM62 based scoring system, we observe that most of the TOP7_full residues were actually favorable, obtaining positive scores. This low conservation is especially relevant considering that intensive studies on TOP7 have revealed the importance of beta-sheet conservation in order to keep its foldability (Boschek et al., 2009; Soares, Boschek, Apiyo, Baird, & Straatsma, 2010; Viana et al., 2013).

**Figure 6 - Supplementary Figure 1.**
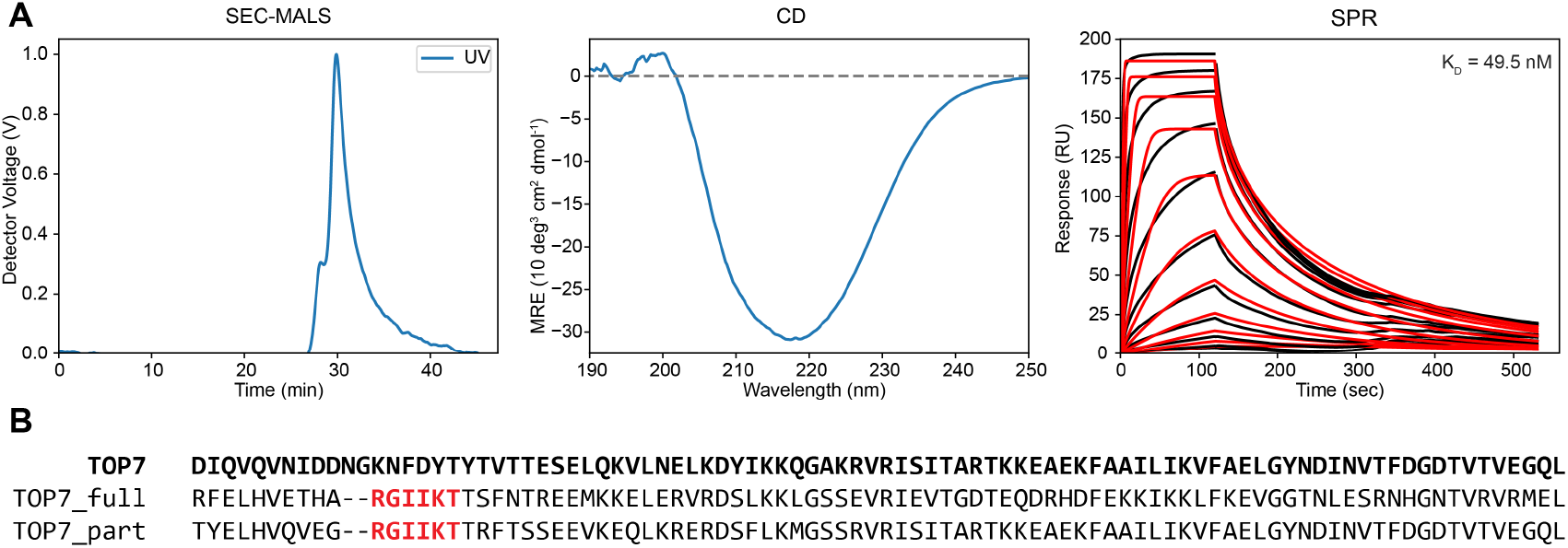
Experimental characterization of TOP7_variants. A) Experimental characterization for the TOP7_partial design: SEC-MALS elution profile (left column); CD wavelength scan spectrum; SPR binding assays with the 101F antibody (right column). Noticeably the TOP7_partia show a CD spectrum notoriously different from WT TOP7 and the TOP7_full design. B) Global sequence alignment of the wild-type protein TOP7 and the computationally designed sequences. Red positions highlight the site IV epitope insertion.

In summary, our results show that FunFolDes was able to repurpose a functionless protein by folding and designing its structure to harbor a functional site, which in this case was a viral epitope. Previously, these computationally designed proteins with embedded viral epitopes were dubbed epitope-scaffolds and showed their biomedical applicability as immunogens that were able to elicit viral neutralizing antibodies(Correia et al., 2014).

## Discussion and Conclusions

The robust computational design of proteins that bear a biochemical function remains an important challenge for present methodologies. The ability to consistently repurpose old folds for new functions or the *de novo* design of functional proteins could bring new insights into the determinants necessary to encode function into proteins (e.g. dynamics, stability, etc.) as well as important advances in translational applications (e.g. biotechnology, biomedical, biomaterials, etc.).

Here, we present the second-generation computational design protocol Rosetta FunFolDes, which was conceived to embed functional motifs into protein topologies, allowing for a global retrofitting of the overall protein topology to favorably host the functional motif. FunFolDes has evolved to incorporate two types of constraints to guide the design process: topological and functional. The former entail the fragments to assemble the protein structure and sets of distance constraints that bias the folding trajectories towards a desired topology; and the latter are the structure of the functional motif inserted and the binding target, if used.

We have extensively benchmarked the protocol, leveraging natural structural and sequence variation of proteins within the same fold, as well as deep mutational scanning data for the computationally designed protein BINDI. In our first benchmark, we observed that with FunFolDes we can efficiently bias the sampling towards improved structural and sequence spaces. Protocol features that enable higher quality sampling in design simulations are extremely important. Improved sampling may contribute to solving some of the major limitations in protein design, related to “junk” sampling, where most of the generated designs are not physically realistic, exhibiting obvious flaws according to general principles of protein structure. Importantly, higher quality sampling will likely contribute to improve the success rate of designs that are tested experimentally. The BINDI benchmark allowed us to test FunFolDes in a system with a large amount of experimental data, which included both sequences and structures. Perhaps the most enlightening observation was that designs that were theoretically within a sequence/structure space productive for binding to the target were rather far from the energetic minimum that the protein fold can achieve in the absence of the binding target. Considering that the large majority of the design algorithms are energy “greedy” and the sequence/structure searches are performed with the central objective of finding the global minimum of the energetic landscape, by introducing functional constrains into the simulations, FunFolDes presents an alternative way of designing functional molecules and efficiently skewing the searches towards off-minima regions of the global landscape. We anticipate that such finding will be more relevant for protein scaffolds that need to undergo a large degree of structural adaptation to perform the desired function. If confirmed that this finding is generalized across multiple design problems, it could be an important contribution for the field of computational protein design.

Furthermore, we used FunFolDes to tackle two design challenges and functionalized two proteins with two distinct viral epitopes. These design challenges were devised to test the applicability of FunFolDes. Importantly, in previous applications FFL always used three-helix bundles as design templates, here we diversified the template folds and used an all-helical protein that is not a bundle (1kx8) and a mixed alpha-beta protein (TOP7), clearly showing the applicability to other folds. For the 1kx8 design series, we evaluated the capability of using distant structural templates as starting topologies as a demonstration of how one can use the many naturally occurring protein structures available and repurpose their function even when the initial template and the target structures are quite different. We obtained stable proteins that where recognized by an anti-RSV antibody with high affinity, showing that in this case, we successfully repurposed a distant structural template for a different function, a task for which other computational approaches (Silva, Correia, & Procko, 2016) would have limited applicability. We see this result as an exciting step forward towards using the wealth of the natural structural repertoire for the design of novel functional proteins.

In a last effort, we functionalized a “functionless” fold, based on one of the first *de novo* designed proteins - TOP7. For us, this challenge has important implications in order to understand the design determinants and biochemical consequences of inserting a functional motif into a protein that was mainly optimized for thermodynamic stability. We were successful in functionalizing TOP7 differently than previous published efforts, where TOP7 was mostly used as a carrier protein with functional motifs fused onto loop regions or side chains grafted in the helical regions, while our functional motif was embedded in the beta sheet of the protein template (Boschek et al., 2009; Soares et al., 2010; Viana et al., 2013). Exciting advances in the area of *de novo* protein design are also yielding many new proteins, which could then be functionalized with FunFolDes, highlighting the usefulness of this approach. Interestingly, we observed that the functionalized version of TOP7 showed a dramatic decrease in thermodynamic stability as compared to the parent protein. While this observation can be the result of many different factors, it is compelling to interpret it as the “price of function”, meaning that to harbor function, the TOP7 protein was penalized in terms of stability, which would be consistent with our findings in the BINDI benchmark example.

Recently, there have also been several *de novo* proteins which were designed for functional purposes (Chevalier et al., 2017); however, these efforts were limited to linear motifs that carried the functions, and the functionalization was mainly accomplished by side-chain grafting (Correia et al., 2010; Kulkarni et al., 2015) and relied on screening of a much larger number of designed proteins.

From our perspective, and considering all the technical improvements, FunFolDes has matured to become a valuable resource for the robust functionalization of proteins using computational design. Here, we present a number of important findings provided by the detailed benchmarks performed and used the protocol to functionalize proteins in design tasks which are representative of some of the common challenges that the broad scientific community faces when using computational design approaches.

## Materials and Methods

### Computational protocol description

Rosetta Functional Folding and Design (FunFolDes) is a general approach for grafting functional motifs into protein scaffolds. It’s main purpose is to provide an accessible tool to tackle specifically those cases in which structural similarity between the functional motif and the insertion region is low, thus expanding the pool of structural templates that can be considered useful scaffolds. This objective is achieved by folding the scaffold after motif insertion while keeping the structural motif static. This process allows the scaffold’s conformation to change and properly adapt to the three dimensional restrictions enforced by the functional motif. The pipeline of the protocol (summarized in Figure 1) proceeds as follows:

i. *Selection of the functional motif* A single or multi-segment motif must be selected and provided as an input. In the most common mode of the protocol dihedral angles, side chain identities and conformations are kept fixed throughout the whole protocol.
ii. *Selection of the protein scaffold*. Searches for starting protein scaffolds can be achieved, but are not limited to, RMSD similarity matches to the Protein Data Bank (PDB) (Rose et al., 2017). The ability of FunFolDes to adapt the scaffold to the needs of the motif widens the structural space of what can be considered as a suitable template. Thus, this step requires human intervention and has to be performed outside of the main protocol.
iii. *Generation of fragment databases*. The usage of fragments lies at the core of many Rosetta protocols, particularly those that perform large explorations of the conformational space required for structure prediction and design. The most standard way of assembling fragment sets is to generate sequence-based fragments using the FragmentPicker application (Kim, Blum, Bradley, & Baker, 2009). Despite the usefulness of the sequence-based fragments in typical design and structure prediction problems, FunFolDes-derived designs depend on the structural content of the template rather than its sequence. Thus we implemented the *StructFragmentMover*, a mover that performs on-the-fly fragment picking based on secondary structure, dihedral angles and solvent accessibility, calculated from the template’s structural information. The typical three- and nine residue-long fragment sets are generated from the fragment database included in the Rosetta tools release.
iv. *Generation of constraints*. Residue-pair distance and backbone dihedral angle constraints can be extracted from the protein scaffold to guide the folding process. These constraints may include the full-length protein or focus in specific segments while allowing a wider flexibility in other regions. Although not required, the use of constraints greatly increases the quality of the sampling. The protocol can be also made aware of other constraint types (such as cartesian constraints) by properly modifying the score functions applied to the *ab initio* stage (Simons, Bonneau, Ruczinski, & Baker, 1999).
v. *Construction of the extended pose*. The extended structure is composed of all the segments of the target motif maintain their native backbone conformation and internal rigid body orientation. The scaffold residues are linearly attached to previously defined insertion points. In multi-segment motif scenarios, the construct will present a chain break between each of the motif composing segments. The number of chain-breaks in the pose scales with the number of segments(n) within a motif always resulting in n-1 chain-breaks. Once the extended pose is assembled, it is represented at the centroid level (all side-chain atoms in a single virtual atom) to reduce the computational cost of the simulation.
vi. *Folding the extended pose*. Fragment insertion is performed to accomplish the folding stage. Kinematics of the pose are controlled through the FoldTree (Wang, Bradley, & Baker, 2007), a system to control the propagation of the torsion angles applied to a structure. In single-segment motif structures, the FoldTree starts in the center of the motif and propagates in opposite directions towards the N- and C-terminal of the protein. In multi-segment motifs, in which the pose bears chain-breaks between each pair of motif segments, the FoldTree has a fixed node in the center of each segment and expands towards both sides (**Figure 1**). The chain-breaks in the structure are marked as cut-points, which avoid further propagation of the kinematic movement throughout the polypeptide chain, and are subjected to a score term to promote their spatial proximity. All the nodes of the FoldTree are placed in the motif segments are kept fixed relative to each other in three-dimensional space; this setup allows for the folding of the protein while maintaining the relative position of all the motif segments.
vii. *Inclusion of the binding target*. If a binding target (protein, nucleic acid or small molecule ligand) is provided, a new FoldTree node is added to the closest residue between the first motif segment and each binding element. Similarly to the multi-segment kinematics, this ensures that the rigid-body orientation between the motif and its target is maintained. FunFolDes can handle simulations with both multi-segment and binding targets simultaneously.
viii. *Folding post-processing*. Folding trajectories are considered successful if they generate structures under a user-defined RMSD threshold of the starting scaffold. In case of a multi-segment motif, a preliminary loop closure will be executed to generate a continuous polypeptide chain, and the kinematic setup maintained to avoid segment displacement during the design step. After the folding stage performed at the centroid level, full atom information is recovered. All the steps necessary to perform the setup of the extended pose (kinematic setup, folding, post-processing) are carried out by a newly implemented mover called *NubInitioMover*.
ix. *Protein design and conformational relaxation*. The folded structure is subjected to iterative cycles of sequence design (Hu, Wang, Ke, & Kuhlman, 2007) and structural relaxation (Tyka et al., 2011) in which the sequence search is coupled with confined conformational sampling (Kuhlman & Baker, 2004). A MoveMap is defined to control backbone dihedrals and side chain conformations of the motif segments and the binding target while allowing for backbone and side-chain exploration of the movable residues. TaskOperations are used to avoid undesired mutations in the functional motif.
x. *Loop closure*. If multi-segment motifs are used, a final loop closure step is required in order to obtain a polypeptide chain without breaks. The *NubInitioLoopClosureMover* performs this last step using the Cyclic Coordinate Descend (CCD) protocol (Wang et al., 2007), while ensuring that the original conformation and rigid-body orientation of the motifs is maintained. After the closure of each cut-point, a final round of fixed backbone design is performed on the residues of the cut-points and surroundings.
xi. *Selection, scoring and ranking*. Finally, the decoys are ranked and selected according to Rosetta energy, structural metrics (core packing, buried unsatisfied polar atoms, etc) (Alford et al., 2017), sequence-based predictions such as secondary structure propensity (Jones, 1999) and folding propensity (Simons, Bonneau, et al., 1999) or any other metrics accessible through RosettaScripts (RS).

The pipeline components described here represent the most standardized version of the FunFolDes protocol. By means of its integration in RS, different stages can be added, removed or modified to tailor the protocol to the specific needs of the design problem at hand.

### Capturing conformational and sequence changes in small protein domains

To test the ability of FunFolDes to recover the required conformational changes to stabilize a given structural motif, we created a benchmark of 14 target cases of proteins with less than 100 residues, named T01 to T14. Each target case was composed of two structures of the same CATH superfamily(Dawson et al., 2017). One of the structures was representative of the shared structural features of the CATH family; we called this structure the reference. The second protein within each target case presents two structural variations with respect to the reference: I) an insertion or deletion (indel) region and II) a conformational change. Direct structural contacts between these two regions make it so that the indel region is the cause for the conformational change. We called this second structure the target (**Figure 2, Table 1**).

**Table 1.** Targets included in the conformational and sequence recovery benchmark. For each of the benchmark target is indicated the CATH superfamily and representatives used in the simulations. (#) indicates the number of segments in the target protein that are considered motif. Motif range indicates the residues considered motif according to the PDB numbering.

| ID | CATH | # | reference | target | motif range |
| --- | --- | --- | --- | --- | --- |
| **T01** | CATH.3.40.140.10 | 1 | 1pgxA | 2pw9C | 69-73 |
| **T02** | CATH.3.30.310.50 | 1 | 3i3wA | 4bjuA | 464-486 |
| **T03** | CATH.3.30.70.980 | 1 | 1lfpA | 1mw7A | 140-150 |
| **T04** | CATH.3.30.70.100 | 1 | 1rjjA | 1lq9A | 19-45 |
| **T05** | CATH.3.10.20.30 | 1 | 2q5wD | 2pkoA | 49-64 |
| **T06** | CATH.2.30.29.30 | 1 | 1c1yB | 1h4rA | 39-59 |
| **T07** | CATH.3.10.20.90 | 1 | 2bkfA | 2al6B | 115-119 |
| **T08** | CATH.3.10.20.90 | 1 | 1wj4a | 1wiaA | 181-200 |
| **T09** | CATH.3.10.20.90 | 1 | 3ny5B | 3phxB | 100-121 |
| **T10** | CATH.3.10.20.310 | 1 | 2x8xX | 2qdfA | 103-121 |
| **T11** | CATH.3.10.320.10 | 1 | 4p5mA | 2bc4C | 56-66 |
| **T12** | CATH.2.40.40.20 | 1 | 1cr5B | 2pjhB | 119-142 |
| **T13** | CATH.2.40.40.20 | 2 | 1cr5B | 2pjhB | 119-142, 168-173 |
| **T14** | CATH.3.30.110.40 | 1 | 1jdqA | 3lvjC | 14-37 |

For each template protein we generated approximately 10000 decoys with FunFolDes by folding the target with the following conditions: 1) the indel region was considered as the motif, meaning that its structural conformation was kept fixed and no mutations allowed; 2) residue-pair distance constraints were derived from the secondary structure elements conserved between reference and the target (constrained region); 3) the region of the protein which showed the largest structural variations (query region) was constraint-free throughout the simulation.

FunFolDes simulations were compared with constrained *ab initio (cst-ab initio)*simulations, the key difference between them being that the cst-*ab initio* simulations allowed for backbone flexibility in the motif region. The comparison between both approaches provides insights on the effects of a static segment in the folding trajectory of the polypeptide chain. In both scenarios a threshold was set after the folding stage where only decoys that had less than 5 Å RMSD from the template were carried to the design stage.

The importance of the input fragments was assessed with our benchmark. Both protocols were tested with sequence-based fragments and structure-based fragments generated on-the-fly by FunFolDes. Comparison between the two types of fragments provides insight into how to utilize FunFolDes in the most productive manner.

Structural recovery was evaluated by RMSD with the target structure. Global RMSD, understood as the minimum possible RMSD given the most optimal structural alignment, was used to assess the overall structural recovery of each decoy population. Local RMSD, was evaluated for the unconstrained (query) region and the motif by aligning each decoy to the template through the constrained segments (excluding the motif). This metric aimed to capture the specific conformational changes required to accommodate the motif into the structure (**Figure 2B, Figure 2 - Supplementary Figure S1B**).

Sequence recovery was evaluated through two different criteria, sequence associated statistics and Hidden Markov Model (HMM) (Eddy, 2011). For the sequence associated statistics, we quantified sequence identity and similarity according to BLOSUM62 for the core residues of each protein, as defined by Rosetta’s *LayerSelector* (Koga et al., 2012). Motif residues, that were not allowed to mutate, were excluded from the statistics. In the second criteria, position specific scoring matrices with inter-position dependency known as Hidden Markov Model (HMM) were used to evaluate fold specific sequence signatures. In this case, the closest HMM to the template structure provided by CATH was used to query the decoys and identify those that matched the HMM under two conditions: I) an e-value under 10 and II) a sequence coverage over 50%. Although these conditions are wide, they were within the variability found between members of CATH superfamilies with high structural and sequence variability like the ones used in the benchmark.

### Target-biased design of protein binders

To assess the performance of FunFolDes in the presence of a binder target we recreated the design of BINDI as a binder for BHRF1 (Procko et al., 2014), the BHRF1 binding motif from the BIM BH3 protein (PDB ID:2WH6 (Kvansakul et al., 2010)) was inserted into a previously described 3-helix bundle scaffold (PDB ID:3LHP (Correia et al., 2010)).

On that account, four different design simulations were performed: one without the binder (no_target) and three in the presence of the binder (static, pack and packmin). The difference between the last three relates to how the binding target was handled. In the static simulations the binding target was kept fixed and no conformational movement in the side chains was allowed throughout the protocol. In the pack simulations the side chains of the binding target were repacked during the binder design stage. Finally, in the packmin simulations the binding target side-chains were allowed to repack and both side-chains and backbone were subjected to minimization. These three target configurations are easily obtained by altering MoveMap definitions, demonstrating the flexibility of the protocol to include variable conditions. In all cases, the two terminal residues on each termini of the binding motif were allowed backbone movement to optimize the insertion in the 3-helix bundle scaffold. For each one of these scenarios, approximately 20000 decoys were generated.

For the no_target simulations the FunFolDes designs were docked to BHRF1 using the inserted motif as guide to assess their complementary and interface metrics. In all the simulations, a final round of global minimization was performed where both proteins of the complex were allowed backbone flexibility. During this minimization, the jump between the design and target was kept fixed to maintain the binding motif and target in place. The final ΔΔG of the complexes was measured after the minimization step to enable comparasions between the no_target decoys and the remaining simulation modes. Structural changes related to this minimization step were evaluated as the global RMSD between each structure before and after the process, this measure is referred to as RMSD drift.

Structural evaluation includes global RMSD against the BINDI design crystal structure (PDB ID: 4OYD (Procko et al., 2014)) as well as local RMSD against regions of interest of BINDI. For the Local-RMSD the structures were aligned through the inserted motif, as it was kept throughout all simulations and with respect to BINDI. The local RMSD analysis was performed over all the helical segments contained in the structures (all H), which provides a measurement of the structural shifts on the secondary structure regions of the designs.

To evaluate the sequence recovery of our simulations we leveraged BINDI’s saturation mutagenesis data analyzed by deep sequencing performed in the by Procko et al (Procko et al., 2014). The experimental fitness of each mutation was summarized in a score matrix where a score was assigned for each amino-acid substitution for the 116 positions of the protein (**Figure 4 - Supplementary Figure 1**). In summary, point mutations that improved BINDI’s binding to BHRF1 are assigned positive scores while deleterious mutations present negative values. These scores are computed based on experimental data where the relative populations of each mutant were compared between a positive population of cells displaying the designs (binders) and negative populations (mutants that display but don’t bind), these experiments have been described in detail elsewhere(Procko et al., 2014). After normalization by the score of the final BINDI sequence in each position, a position sequence specific matrix (PSSM) was created. Like the original data, this matrix also assigns a positive score to each point mutation if it resulted in an improved binding for the design. This normalization provides a score of 0 for the BINDI sequence, which is useful as a reference score.

### Repurposing naturally occurring folds for a new functions

To experimentally validate the capabilities of FunFolDes and insert functional sites in structurally distant templates, we decided to transfer the site II epitope from the Respiratory Syncytial Virus (RSV) protein F (PDB ID:3IXT (McLellan, Chen, Kim, et al., 2010)) into heterologous scaffolds. This is a continuous, single segment, helix-loop-helix conformation epitope. The main objective was to challenge the capabilities of FunFolDes to reshape the structure of the scaffold to the requirements of the functional motif, we aimed to search for insertion segments with RMSDs towards the site II structure higher than 2 Å.

We searched for host scaffolds using MASTER (Zhou & Grigoryan, 2015) where we used the full-length site II segment as a query against a subset of 17539 protein structures from the PDB composed of 30% non-redundant sequences included in the MASTER distribution. The RMSD between the query and segments on the scaffolds were assessed using backbone C_α_s. All matches with RMSD_Cα_ < 5.5 Å relative to site II were recovered and further filtered by protein size, where only proteins between 50 and 100 residues were kept. These scaffolds were then ranked regarding antibody-binding compatibility, where each match was realigned to the antibody-epitope complex and steric clashes between all glycine versions of the scaffold and antibody were quantified using Rosetta. All matching scaffolds with ΔΔG values above 100 REU were discarded under the assumption that their compatibility with the antibody binding mode was to low. The remaining scaffolds were visually inspected and PDB ID: 1kx8 (Lartigue et al., 2002) (RMSDeα = 2.37 Å) was selected for design with FunFolDes.

The twenty-one residues from the site II epitope (motif) as present in 3IXT were grafted into a same sized segment (residues 79-100) of 1kx8 using the *NubInitioMover*. Up to three residues in each insertion region of the motif were allowed backbone flexibility in order to allow proper conformational transitions in the insertion points. Atom pair constraints with a standard deviation of 3 Å were defined for all template residues, leaving the motif residues free of constraints. The generous standard deviation was set up to favour necessary conformational changes to allow the optimal fitting of the motif within the topology. Regardless, the total allowed deviation for template was limited at 5 Å to ensure the retrieval of the same topology. In this design series we used sequence-based fragments generated with the 1kx8 native sequence. Three cycles of design/relax were performed on the template residues with the *FastDesignMover*.

A first generation of 12500 designs was ranked according to Rosetta energy. From the top 50 decoys, only one presented the motif without distortions on the edges derived from the allowed terminal flexibility. This decoy was used as template on the second generation of FunFolDes to enhance the sampling of properly folded conformations, with the same input conditions as the previous one.

In the second generation, the top 50 decoys according to Rosetta energy were further optimized through additional cycles of design/relax. After a selection based on Rosetta energy, buried unsatisfied polars and secondary structure prediction using PSIPRED (Jones, 1999), a total of 7 designs were manually optimized and selected for experimental characterization. After the initial characterization, designs with added disulphide bridges were generated in order to improve protein stability and affinity (Figure 4 - Supplementary Figures 1 and 2).

### Functionalization of a functionless fold

In a second effort to test the design capabilities of FunFolDes we sought to insert a functional motif in one of the first de novo designed proteins - TOP7 (PDB ID: 1QYS) (Kuhlman et al., Science, 2003).

Six residues from the complex between the antibody 101F and the peptide-epitope, corresponding to residues 429-434 (RGIIKT) on the full-length RSV F protein (McLellan, Chen, Chang, et al., 2010), were grafted into the edge strand of the TOP7 backbone using FunFolDes. The choice between epitope and hosting scaffold was made based on the secondary structure adopted by the epitope and the acceptable structural compatibility of the TOP7 structure, the RMSDCα between the epitope an the insertion segment was 2.07 Å.

To ensure that the majority of the β-strand secondary structure was maintained throughout the grafting protocol, the epitope motif was extended by one residue and a designed 4-residue β-strand (KVTV) pairing with the backbone of the C-terminal epitope residues was co-grafted as a discontinuous segment into the adjacent strand in the TOP7 backbone. With this strategy we circumvented a Rosetta sampling limitation, where often times is necessary an extensive set of constrains to achieve the desired backbone hydrogen-bond pairing on beta-strands (Marcos et al., 2017). After defining the motif consisting of the epitope plus the pairing strand and the sites of insertion on the TOP7 scaffold, FunFolDes was used to graft the motif.

Backbone flexibility was allowed for the terminal residues of the functional motif and a β-turn connection between the two strands was modelled during the folding process (*NubInitioMover*). During the folding process, 101F antibody was added to the simulation in order to limit the explored conformational space towards binding productive designs. Finally, the *NubInitioLoopClosureMover* was applied to ensure that a proper polypeptide chain was modelled and no chain-brakes remained, a total of 800 centroid models were generated after this stage. Next, we applied an RMSD filter to select scaffolds with similar topology to TOP7 (< 1.5 Å) and a hydrogen bond long-range backbone score (HB_LR term) to favour the selection of proteins with proper beta-sheet pairing. The top 100 models according the HB_LR score and that also fulfilled the RMSD filter, were then subjected to an iterative sequence-design relax protocol, alternating fixed backbone side-chain design and backbone relaxation using the *FastDesignMover*. Designable positions were limited to a subset of residues according to their position in the core or surface of the protein and secondary structure identity. Two different design strategies were pursued: I) partial design - amino acid identities of the C-terminal half of the protein (residues 45 through 92) were retained from TOP7 while allowing repacking of the side chains and backbone relaxation; II) full-design - the full sequence space in all residues of the structure (with the exception of the 101F epitope) was explored. No backbone or side chain movements were allowed in the 6-residue epitope segment whereas the adjacently paired β-strand was allowed to both mutate and relax. Tight Cα atom-pair distance constraints (standard deviation of 0.5 Å) were used to restrain movements of the entire sheet throughout the structural relaxation iterations.

From the 100 designs generated, only those that passed a structural filter requiring that 80% of secondary structure composition of the β-sheet after backbone relaxation were selected for further analysis. The 93 designs passing this filter were evaluated based on several metrics such as: REU, hydrogen-bond long-range backbone interactions and core packing. The best-scored designs were finally submitted to human-guided optimisation, mutations of single surface residues (1-3) and shortening of the connecting loop between the two inserted strands using the Rosetta Remodel application.

Interestingly, in an attempt to reproduce the same grafting exercise with *MotifGraftMover* (Silva et al., 2016), this resulted in non-resolvable chain breaks when trying to graft either the two segment-motif or the epitope alone into the TOP7 scaffold.

### Protein Expression and Purification

DNA sequences of the designs were purchased from Twist Bioscience. For bacterial expression the DNA fragments were cloned via Gibson cloning into a pET21b vector containing a C-terminal His-tag and transformed into *E. coli* BL21(DE3). Expression was conducted in Terrific Broth supplemented with ampicillin (100 μg/ml). Cultures were inoculated at an OD_600_ of 0.1 from an overnight culture and incubated at 37°C with a shaking speed of 220 rpm. After reaching OD_600_ of 0.7, expression was induced by the addition of 1 mM IPTG and cells were further incubated for 4-5h at 37°C. Cells were harvested by centrifugation and pellets were resuspended in lysis buffer (50 mM TRIS, pH 7.5, 500 mM NaCl, 5% Glycerol, 1 mg/ml lysozyme, 1 mM PMSF, 1 μg/ml DNase). Resuspended cells were sonicated and clarified by centrifugation. Ni-NTA purification of sterile-filtered (0.22 μm) supernatant was performed using a 1 ml His-Trap™ FF column on an ÄKTA pure system (GE healthcare). Bound proteins were eluted using an imidazole concentration of 300 mM. Concentrated proteins were further purified by size exclusion chromatography on a Superdex™ 75 300/10 or a Superdex™ Hiload 16/600 75 pg column (GE Healthcare) using PBS buffer (pH 7.4) as mobile phase.

For IgG expression, heavy and light chain DNA sequences were cloned separately into pFUSE-CHIg-hG1 (InvivoGen) mammalian expression vectors. Expression plasmids were co-transfected into HEK293-F cells in FreeStyle™ medium (Gibco™) using polyethylenimine (Polysciences) transfection. Supernatants were harvested after 1 week by centrifugation and purified using a 5 ml HiTrap™ Protein A HP column (GE Healthcare). Elution of bound proteins was accomplished using a 0.1 M glycine buffer (pH 2.7) and eluents were immediately neutralized by the addition of 1 M TRIS ethylamine (pH 9). The eluted IgGs were further purified by size exclusion chromatography on a Superdex 200 10/300 GL column (GE Healthcare) in PBS buffer (pH 7.4). Protein concentrations were determined by measuring the absorbance at 280 nm using the sequence calculated extinction coefficient on a Nanodrop (Thermo Scientific).

### Circular Dichroism (CD)

Far-UV circular dichroism spectra of designed scaffolds were collected between a wavelength of 190 nm to 250 nm on a Jasco J-815 CD spectrometer in a 1 mm path-length quartz cuvette. Proteins were dissolved in PBS buffer (pH 7.4) at concentrations between 20 μM and 40 μM. Wavelength spectra were averaged from two scans with a scanning speed of 20 nm min^−1^ and a response time of 0.125 sec. The thermal denaturation curves were collected by measuring the change in ellipticity at 220 nm from 20 to 95°C with 2 or 5 °C increments.

### Size-exclusion Chromatography combined with Multi-Angle Light-Scattering (SEC-MALS)

Multi-angle light scattering was used to assess the monodispersity and molecular weight of the proteins. Samples containing between 50 −100 μg of protein in PBS buffer (pH 7.4) were injected into a Superdex™ 75 300/10 GL column (GE Healthcare) using an HPLC system (Ultimate 3000, Thermo Scientific) at a flow rate of 0.5 ml min^−1^ coupled in-line to a multi-angle light scattering device (miniDAWN TREOS, Wyatt). Static light-scattering signal was recorded from three different scattering angles. The scatter data were analysed by ASTRA software (version 6.1, Wyatt)

### Surface Plasmon Resonance (SPR)

To determine the dissociation constants of the designs to the mota or 101F antibody, surface plasmon resonance was used. Experiments were performed on a Biacore 8K at room temperature with HBS-EP+ running buffer (10 mM HEPES pH 7.4, 150 mM NaCl, 3mM EDTA, 0.005% v/v Surfactant P20) (GE Healthcare). Approximately 1200 response units of mota or 101F antibody were immobilized via amine coupling on the methylcarboxyl dextran surface of a CM5 chip (GE Healthcare). Varying protein concentrations were injected over the surface at a flow rate of 30 μl/min with a contact time of 120 sec and a following dissociation period of 400 sec. Following each injection cycle, ligand regeneration was performed using 3M MgCh (GE Healthcare). Data analysis was performed using 1:1 Langmuir binding kinetic fits within the Biacore evaluation software (GE Healthcare).

### Availability

FunFolDes is available as part of the Rosetta software suite and is fully documented in the Rosetta Commons documentation website as one of the Composite Protocols. All data and scripts necessary to recreate the analysis and design simulations described in this work are available at https://github.com/lpdi-epfl/FunFolDesData.

## Contributions

J.B. coded the algorithm described. A.S coded the *StructFragMover*. K.S., A.B., A.S. and F.S. performed computational design simulations. S.W., K.S, C.Y., A.B., F.S., S.V., R. L., M. V. and S.R. contributed to experimental characterization of the designed proteins. J.B. and B.E.C. designed the study and wrote manuscript.

## Acknowledgements

We would like to acknowledge the High performance computing facility (SCITAS) for their technical support. We would also like to acknowledge the Swiss National Supercomputing Centre (CSCS) for their support in computing time. We would like to thank the protein expression and characterization platform (PCRYCF/PECF) for their support with mammalian expression and access to analytical instrumentation. We would like to thank Erik Procko for providing the data for the saturation mutagenesis on BINDI.

